# ISWI catalyzes nucleosome sliding in condensed nucleosome arrays

**DOI:** 10.1101/2023.12.04.569516

**Authors:** Petra Vizjak, Dieter Kamp, Nicola Hepp, Alessandro Scacchetti, Mariano Gonzalez Pisfil, Joseph Bartho, Mario Halic, Peter B. Becker, Michaela Smolle, Johannes Stigler, Felix Mueller-Planitz

## Abstract

How chromatin enzymes work in condensed chromatin and how they maintain diffusional mobility inside remains unexplored. We investigated these challenges using the *Drosophila* ISWI remodeling ATPase, which slides nucleosomes along DNA. Folding of chromatin fibers did not affect sliding *in vitro*. Catalytic rates were also comparable in- and outside of chromatin condensates. ISWI cross-links and thereby stiffens condensates, except when ATP hydrolysis is possible. Active hydrolysis is also required for ISWI’s mobility in condensates. Energy from ATP hydrolysis therefore fuels ISWI’s diffusion through chromatin and prevents ISWI from cross-linking chromatin. Molecular dynamics simulations of a ‘monkey-bar’ model in which ISWI grabs onto neighboring nucleosomes, then withdraws from one before rebinding another in an ATP hydrolysis-dependent manner qualitatively agree with our data. We speculate that ‘monkey-bar’ mechanisms could be shared with other chromatin factors and that changes in chromatin dynamics caused by mutations in remodelers could contribute to pathologies.

## Introduction

Eukaryotes prevent non-specific, cation-induced aggregation of genomic DNA by wrapping their DNA around histones, thereby forming nucleosomes (Burak et al., 2003; Post & Zimm, 1982). Nucleosomes allow for regulated and reversible condensation of genomic DNA and have evolved as an important medium for epigenetic gene regulation.

Nucleosomal DNA can fold into various higher order structures. In low ionic strength environments, chromatin forms an extended structure referred to as the 10-nm fiber (Olins & Olins, 1974; Woodcock, 1973). Moderate ionic strength promotes folding into different but defined higher order structures, collectively referred to as ’30-nm fibers’ (Finch & Klug, 1976). Cellular chromatin, however, appears to assume variable, irregular structures (Ou et al., 2017). It may contain interdigitated 10-nm fibers (Adhireksan et al., 2020; Maeshima et al., 2016) and small folded nodules of chromatin, including tri- and tetranucleosomes (Hsieh et al., 2015; Ricci et al., 2015).

Recently, it was discovered that reconstituted chromatin can undergo phase separation (Gibson et al., 2019; Strickfaden et al., 2020). Phase separation is a property of multivalent polymers incl. many proteins and nucleic acids. They can demix from the solution into polymer-rich condensates and a polymer-depleted solution. Biomolecular condensates harbor distinct chemical environments compared to their surrounding solution. They could, for instance, be enriched or depleted for cellular enzymes or metabolites, providing an additional regulatory layer of cellular processes (Y. Zhang et al., 2021).

Concentrations can massively increase upon condensate formation. For example, sub-micromolar concentrations of soluble nucleosomes form salt-induced condensates that contain ∼0.34 mM nucleosomes (Gibson et al., 2019). Such a high concentration rivals nucleosome concentrations observed in highly condensed metaphase chromosomes *in vivo* (0.52 mM) (Hihara et al., 2012).

Most nuclear processes, including transcription and replication, are thought to be inhibited by the presence of nucleosomes (Kornberg & Lorch, 2020). Intuitively, one would expect that folding of chromatin into various higher-order structures and formation of chromatin condensates could pose additional barriers for nuclear factors. For instance, access to specific DNA sequences may be restricted if they are buried deep inside a folded structure, nucleosomal epitopes may become unavailable, enzymes may be excluded from condensates, and even if they are not, diffusion through dense nucleosome fibers may become slow and inefficient. Remarkably, many chromatin enzymes are mobile in the nucleus (Kim et al., 2021). Since the high concentration of nucleosomes in the nucleus well exceeds the dissociation constants (*K*_D_) of most chromatin-associated enzymes, mechanisms must exist that prevent quantitative binding of the enzymes to chromatin and allow sufficient diffusional mobility.

We address these and other fundamental questions using *Drosophila melanogaster* ISWI as a prototypical nucleosome remodeling enzyme. ISWI slides nucleosomes along DNA in an ATP hydrolysis-dependent fashion (Längst et al., 1999; Hamiche et al., 1999). It can slide internal nucleosomes in reconstituted nucleosome arrays (Klinker, Mueller-Planitz, et al., 2014; Ludwigsen et al., 2018; Mueller-Planitz et al., 2013; Schram et al., 2015). However, if and how fiber folding and condensation have an impact on nucleosome remodeling has not been systematically investigated, despite the physiological importance (Boyer et al., 2000; Logie et al., 1999). A dense and folded environment could conceivably obstruct remodeling activity, or possibly enhance it as observed for other enzymes (Peeples & Rosen, 2021).

## Results

### Intramolecular chromatin folding does not impede remodeling

We reconstituted nucleosomes on DNA with 25 repeats of the Widom ‘601’ nucleosome positioning sequence (Fig. 1a). 13 of these repeats contain a unique restriction site, such as a BamHI site, which becomes exposed during nucleosome remodeling (Ludwigsen et al., 2018; Mueller-Planitz et al., 2013). By measuring accessibility with BamHI, we compared remodeling of folded and unfolded chromatin arrays.

**Figure 1.**
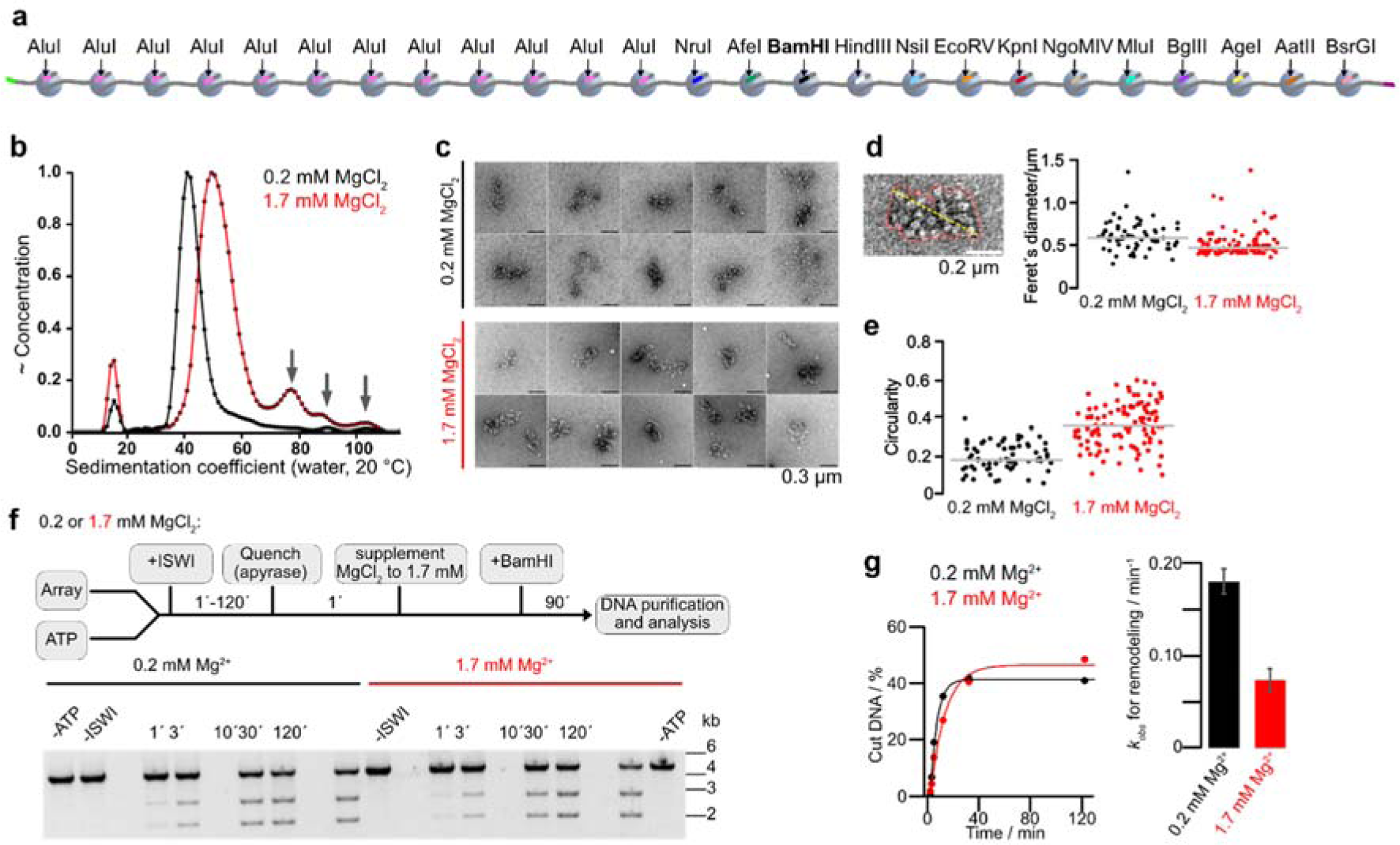
Intramolecular chromatin fiber folding does not impede remodeling. **a**, A 25mer nucleosome array assembled on 25 repeats of 197 bp long Widom-601 DNA derivatives. 13 repeats contain unique restriction enzymes sites. **b**, Sedimentation velocity analysis of 25mer arrays in a buffer supplemented with 0.2 mM (black; peak at 42.4 S) or 1.7 mM MgCl2 (red; peak at 51.3 S). The larger S value at the higher Mg^2+^-concentration indicated stronger compaction of arrays without substantial oligomer formation (arrows). **c**, Representative micrographs of 25mer arrays from negative stain electron microscopy dissolved in a buffer supplemented with indicated MgCl2 concentrations. **d**, **e,** Single-particle analysis of negative stain EM micrographs. N = 67 (0.2 mM MgCl2); N = 107 (1.7 mM MgCl2). Outlines of single particles (red line) were determined with a trainable Weka segmentation in ImageJ. Feret’s diameter (the maximum distance between two parallel tangential lines, yellow line) and circularity were calculated for all outlines. **f**, Top: Schematic of the BamHI accessibility assay. Bottom: Gel analysis of remodeling time courses. Reactions contained 4 nM arrays and 5 µM Mg-ATP and were started by addition of 200 nM ISWI. **g**, Left: quantification of gel in *f* and exponential fits of the time courses. Right: rate coefficients from single exponential fits. Bars are mean values of two independent experiments, error bars their minimal and maximal values.

After *in vitro* nucleosome reconstitution, three independent quality controls confirmed saturation of array DNA with nucleosomes (Fig. S1a). Analytical ultracentrifugation and negative stain electron microscopy showed intramolecular folding of chromatin arrays but no substantial multimerization at Mg^2+^ concentrations of 1.7 mM. (Fig. 1b, c). A lower Mg^2+^ concentration (0.2 mM) served as a loosely folded control. Whereas single particles exhibited structural heterogeneity in both conditions, their Feret’s diameter decreased, and their circularity increased with increasing Mg^2+^ concentration, consistent with Mg^2+^-induced intramolecular folding (Fig. 1d, e).

Induction of fiber folding by supplementation of Mg^2+^ to 1.7 mM reduced rate constants for remodeling by 2.5±0.2 fold (Fig. 1f, g). Folding however is not the major cause of this drop in activity, as the higher Mg^2+^ concentration also reduced the ATPase activity in presence of non-folding mononucleosomes (Fig. S1b).

Internal nucleosomes in nucleosome fibers are more stable and less accessible than external, end-positioned nucleosomes (Hagerman et al., 2009; Poirier et al., 2008, 2009; Schram et al., 2015). To confirm that the effect we observe is not dependent on the linear nucleosome position in an array, we cloned a second version of the 25-mer array, in which the positions of the restriction sites are reversed (Fig. S1c). Supporting our earlier modeling results (Schram et al., 2015), ISWI remodeled nucleosomes with identical rate constants regardless of their location in the array (compare Fig. S1d, e with Fig. 1f, g). We conclude that Mg^2+^-induced intramolecular folding of the chromatin fiber does not impede nucleosome remodeling by ISWI.

### ISWI slides nucleosomes inside chromatin condensates

Mg^2+^ also promotes intermolecular interactions between array molecules at elevated array concentrations (Maeshima et al., 2016). Spherical condensates formed upon addition of physiologically relevant concentrations of Mg^2+^ (1 mM and 5 mM Mg^2+^; Fig. 2a) (Gibson et al., 2019; Strickfaden et al., 2020). The physical nature of these condensates can range from liquid-to solid-like depending on conditions and nucleosome array length (Gibson et al., 2019, 2023; Strickfaden et al., 2020).

**Figure 2.**
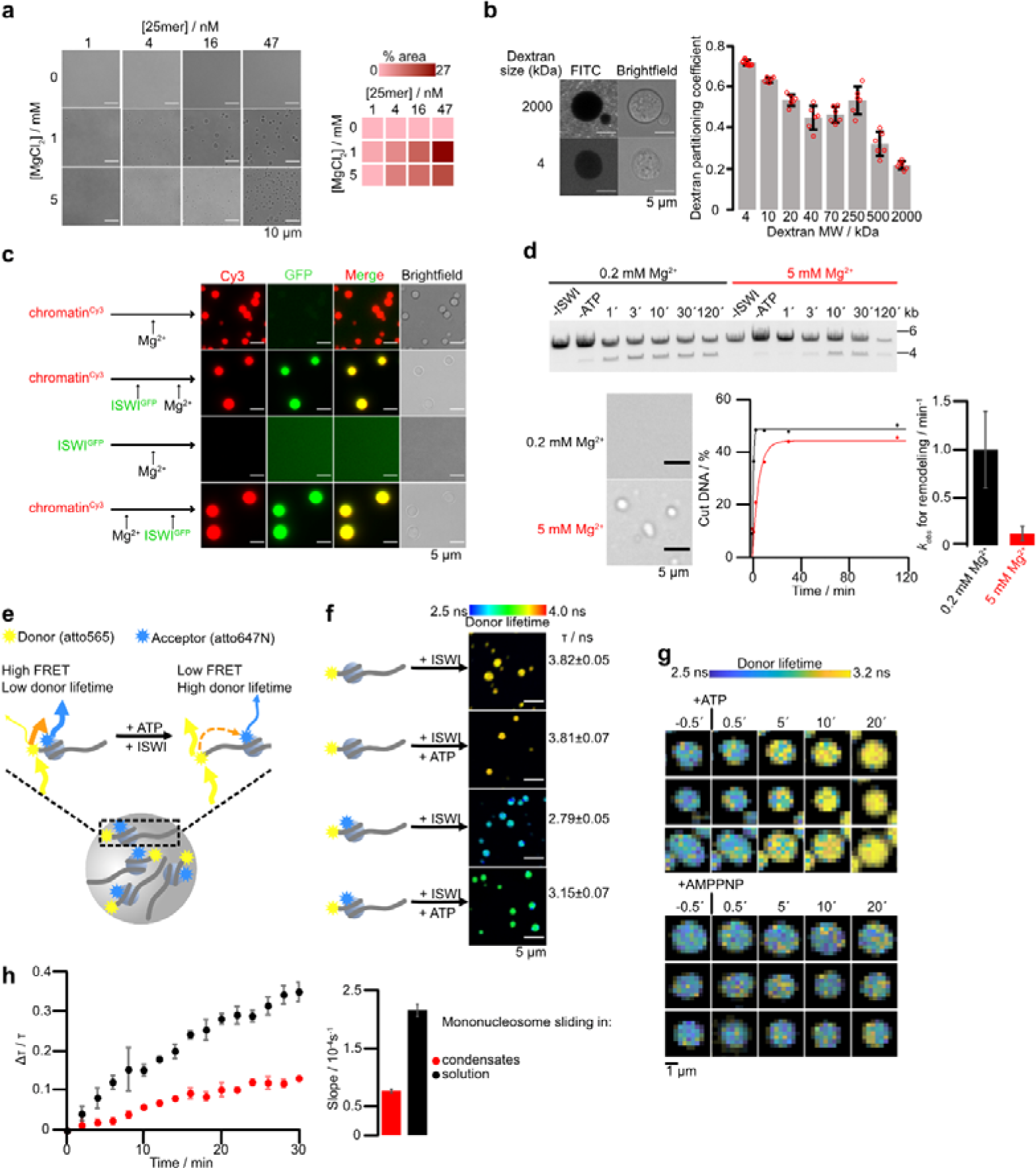
ISWI slides nucleosomes inside chromatin condensates. **a**, Condensate formation depends on nucleosome and MgCl2 concentration. Left: Condensate formation for different MgCl2 and 25mer array concentrations. Right: Percentage of the field of view occupied by condensates. **b**, Partition coefficient of FITC-labeled dextrans of indicated molecular weights. Bars are averages from six condensates for each dextran, errors are SD. **c**, Colocalization experiments with GFP-ISWI (1.1 µM) and Cy3-labeled condensates (0.05 µM 25mers). Condensates were induced by addition Mg^2+^ (5 mM) either before or after addition of GFP-ISWI. **d,** Nucleosome sliding time courses (top and bottom left) in absence and presence of condensates (bottom right). Sliding was measured by KpnI accessibility of 25mer arrays (15 nM) with 1 mM Mg-ATP. Reactions were started by addition of 750 nM ISWI. Bars are mean values of two independent experiments, error bars their minimal and maximal values. **e,** Principle of the FLIM-FRET sliding assay. FRET nucleosomes with a donor dye (atto565) coupled to H2A K119C and an acceptor dye (atto647) attached to the octamer-proximal end of 207 bp Widom-601 DNA were spiked to array condensates. Upon nucleosome sliding, the FRET efficiency decreases, prolonging the donor fluorescence lifetime. **f,** The donor lifetime of FRET-nucleosomes in condensates increased after addition of ISWI and Mg-ATP but not in negative controls (donor-only nucleosomes, or without Mg-ATP). One out of two independent replicates with similar results is shown. Bars are averages and error bars are SD of lifetimes across ten fields of view of the same sample, with each field of view imaged 30 times, totaling 300 images per condition. Reactions contained 45 nM 25mers, 125 nM FRET nucleosomes, 625 nM ISWI and were imaged four hours after addition of Mg-ATP. **g,** FLIM time lapse microscopy. Mg-ATP or Mg-AMPPNP (both 1 mM) were flown into the imaging chamber at t = 0. Individual condensates are pictured. One of two independent replicates with similar results is displayed. **h,** Left: FLIM-FRET time courses in condensates (N=3) and solution (N=2). Points are mean values of independent experiments, error bars their minimal and maximal values. Condensate reactions contained 45 nM 25mers, 125 nM FRET nucleosomes, and 625 nM ISWI and were started with 1 mM Mg-ATP. Solution reactions contained 1125 nM unlabeled 0N60 nucleosomes instead of arrays. Right: Initial velocities of time courses obtained from linear fits of the time courses. Bars are mean values of independent experiments, error bars their minimal and maximal values.

Given their high nucleosome concentrations, these condensates could potentially exclude large protein complexes. To determine the size exclusion limit of the condensates, we used a series of fluorophore-labeled dextrans of different molecular weights and determined their partition coefficients (Fig. 2b). Dextrans were progressively excluded from chromatin with increasing molecular mass. Nevertheless, even 500 kDa dextrans, which possess a Stokes radius (15.1 nm) (Goins et al., 2008) equivalent to a 6 MDa protein complex, were able to enter the condensates. The data suggest that the chromatin condensates are porous or pliable enough to accommodate large particles.

Our dextran results encouraged us to evaluate whether ISWI can also penetrate condensates. We premixed GFP-labeled ISWI with Cy3-labeled chromatin arrays in the absence of nucleotides and induced phase separation by addition of Mg^2+^. ISWI-GFP was strongly enriched in condensates (Fig. 2c). Control experiments proved that GFP or ISWI-GFP did not form condensates by themselves and that GFP on its own does not enrich in chromatin condensates (Fig. 2c and Fig. S2a). Importantly, order of addition experiments showed that ISWI-GFP could penetrate condensates even after their formation with similar efficiency (Fig. 2c). ISWI reached a concentration of ∼4 µM in condensates (Fig. S2b).

The intermolecular interactions between nucleosomes in condensates could conceivably prevent ISWI-mediated sliding of nucleosomes. Using the restriction enzyme accessibility assay, we clearly detected remodeling under condensate conditions (5 mM Mg^2+^; Fig. 2d and Fig. S2c). The observed rate constant for sliding was reduced about eight-fold at the higher Mg^2+^ concentration. Condensation, however, is not the major cause of this drop in the activity, as the higher Mg^2+^ concentration also reduced the ATPase activity in presence of mononucleosomes to a similar extent (Fig. S1b).

To test if remodeling takes place inside of condensates, we developed an imaging-based nucleosome sliding assay (Fig. 2e). We employed double-fluorophore labeled mononucleosomes that can report on nucleosome sliding via changes in FRET (Yang & Narlikar, 2007). These FRET nucleosomes readily partitioned into condensates (Fig. S2d). FRAP of individual condensates showed little exchange of the FRET nucleosomes between the condensate and the surrounding dilute phase (Fig. S2e). Fluorescence lifetime imaging (FLIM) then allowed us to image FRET levels in- and outside of condensates. The donor lifetime of FRET nucleosomes was quenched compared to nucleosomes that were only labeled with the donor fluorophore or whose acceptor fluorophore was bleached, confirming FRET (Fig. 2f and Fig. S2f, g). Importantly, addition of ISWI and Mg-ATP to FRET nucleosomes, but not to donor-only nucleosomes, increased the donor lifetime, indicative of nucleosome sliding (Fig. 2f and Fig. S2f). Time course measurements after flowing in Mg-ATP to ISWI-containing condensates reveal a steady increase in the fluorescence lifetime (Fig. 2g, Movie S1). No lifetime changes were recordable after addition of the non-hydrolyzable ATP analog Mg-AMPPNP as a negative control (Movie S2). We conclude that FLIM-FRET is a sensitive technology to visualize nucleosome sliding with spatial resolution.

To quantify if and to what extent condensates pose a barrier for remodeling, we determined the initial velocities for nucleosome sliding inside condensates (0.77±0.06 s^-1^) and compared it to remodeling in solution (2.15±0.15 s^-1^). We obtained the latter by using mononucleosomes that do not phase separate under these conditions at equivalent bulk concentrations (1.2 µM) under otherwise identical conditions (Fig. 2h). Use of five times higher Mg-ATP concentrations gave similar results (Fig. S2h), arguing that Mg-ATP was saturating both in solution and condensates. ISWI concentrations inside condensates were substoichiometric, reaching 4.7 ± 0.6 µM (Fig. S2b), which is well below the nucleosome concentration in condensates (225 ± 59 µM; see below). In summary, condensates slowed down remodeling by 2.6-fold, suggesting that condensation poses no substantial barrier for remodeling; instead it may fine tune the remodeling activity.

### ISWI requires active hydrolysis to diffuse through condensates

Nucleosome concentrations inside of condensates reach remarkably high levels, 225 ± 59 µM, equivalent to ∼45 g/l, as determined by holotomography (Fig. 3a), a label-free methodology that can determine concentrations from the refractive index. Our measurement using 25-mer arrays agreed well with previous estimates that employed 4- and 12-mer nucleosome arrays (195 to 550 µM) using electron tomography and fluorescence microscopy (Gibson et al., 2019; M. Zhang et al., 2022). A broad range of conditions (array length and buffer conditions) thus leads to relatively static nucleosome densities in condensates. Of note, *in vivo* nucleosome concentrations fall into the same regime (100 µM to 500 µM) (Hihara et al., 2012; Weidemann et al., 2003), making *in vitro* condensates a model for studying challenges encountered by chromatin enzymes in the crowded nucleus.

**Figure 3.**
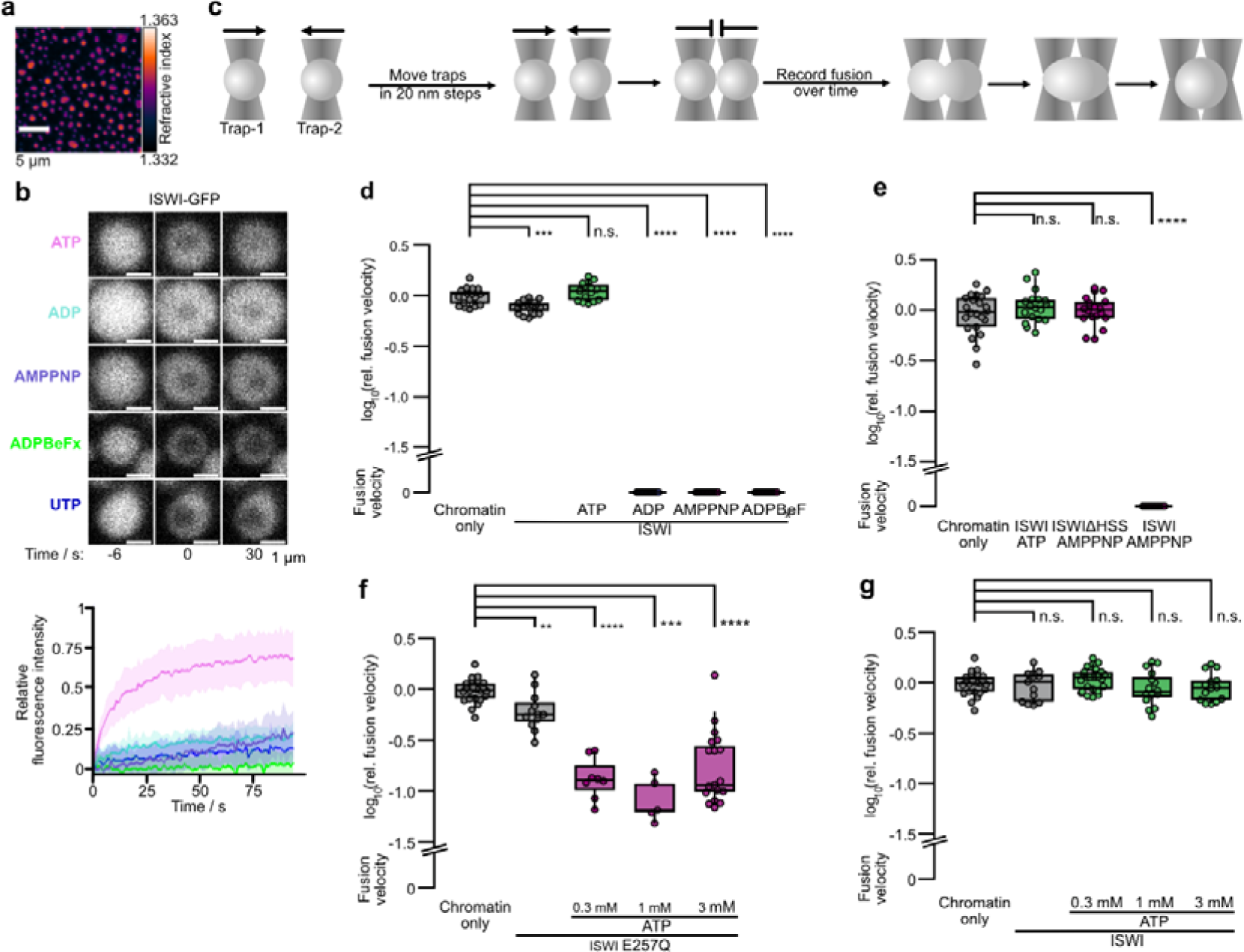
ATP hydrolysis powers mobility of ISWI and prevents ISWI-mediated hardening of condensates. **a**, Holotomogram of chromatin condensates. The average nucleosome concentration (225 ± 59 µM) was determined from the refractive index. One of two independent replicates is depicted. **b,** Left: Partial-condensate FRAP of ISWI-GFP in presence of indicated nucleotides (all 0.77 mM). Only the addition of ATP allowed a fast recovery. Bottom: Quantification of FRAP time courses. Lines are averages, shaded areas SD of 20 condensates for AMPPNP and UTP, 15 for ADP, 25 for ADP-BeFx and 30 for ATP. **c,** Schematic of condensate fusion experiments with optical tweezers. **d,** Fusion velocities of condensates measured by optical trapping. ISWI in presence of non-hydrolysable nucleotides but not of ATP, slowed down fusion. Chromatin-only and nucleotide-free ISWI conditions were replicated three times. **e,** AMPPNP did not slow down fusion when ISWI lacked its HSS domain. **f,** Addition of ATP to an ATPase dead mutant of ISWI (E257Q) slowed down condensate fusion. **g,** Addition of ATP to wild type ISWI did not affect condensate fusion. Data in d–g are log10-fold changes in velocities relative to the mean of the chromatin only condition. Unless stated otherwise, all data were independently replicated twice per condition.

At such a high concentration, well above the dissociation constant of ISWI (<1 µM; (Leonard & Narlikar, 2015)), the enzyme would be expected to be constantly bound to nucleosomes, presumably limiting its mobility. Yet, we found above that condensates do not abrogate remodeling, suggesting a certain mobility of the enzyme.

To probe ISWÍs dynamics inside condensates, we performed FRAP of GFP-labeled ISWI. After bleaching the central area of condensates, there was a poor recovery in the absence of nucleotides and with saturating concentrations of Mg-ADP or Mg-AMPPNP (Fig. 3b and Fig. S3a), consistent with the expectation that ISWI is highly immobile in condensates. Strikingly, however, Mg-ATP addition rendered ISWI strongly dynamic, suggesting that active ATP hydrolysis is required not only for productive remodeling, but also to ensure rapid diffusion of ISWI within condensates.

### ATP hydrolysis prevents ISWI-mediated hardening of condensates

An intriguing model to explain the mobility of ISWI through condensates in the presence of ATP would be that chromatin remodeling by ISWI makes the condensate more fluid. ISWI would essentially act as a ‘molecular stir bar’ (Larson & Narlikar, 2018). However, fluorophore-labeled nucleosome arrays were not mobile, as assessed by FRAP, even after prolonged incubation with ISWI and Mg-ATP (Fig. S3b). We conclude that enhanced fluidity of chromatin upon remodeling is unlikely to explain efficient diffusion of ISWI in presence of ATP.

ISWI has two known DNA binding domains, its ATPase domain and its HSS domain (Grüne et al., 2003), allowing it to bind two nucleosomes (Bhardwaj et al., 2020; L. Li et al., 2023; Yamada et al., 2011) potentially on neighboring DNA fibers. ISWI-mediated bridges between neighboring fibers would alter mechanical properties of condensates. To test this prediction, we employed optical tweezers to fuse condensates in a controlled manner (Fig. 3c) and recorded the fusion velocity (Fig. S3c), which depends on the viscoelastic properties of condensates (Wang et al., 2018). The number of fusion events decreased with the length of the nucleosome array, consistent with a recent report (Muzzopappa et al., 2021), and with increased Mg^2+^ concentration (Fig. S3d). Higher ISWI concentrations also slowed down the fusion velocity (Fig. S3e), which, at least partially, can be explained by an increase in viscosity (Fig. S3f). For these reasons, the following experiments were performed with 13mer arrays in 1 mM MgCl2 and substoichiometric ISWI amounts, typically 1:5 ISWI to nucleosomes.

Compared to nucleotide-free ISWI, the fusion velocities drastically slowed down when we added Mg-AMPPNP, Mg-ADP-BeFx or Mg-ADP to ISWI-containing condensates (Fig. 3d, Movie S3). The presence of ISWI’s HSS domain was required for this effect (Fig. 3e). Fusion velocities also shrank when we added Mg-ATP to an ATP hydrolysis-defective ISWI mutant (ISWIE257Q; Fig. 3f). Strikingly, however, Mg-ATP did not slow down fusion when we used wild-type ISWI instead (Fig. 3g, Movie S4). In control experiments, we ruled out that nucleotides or excess Mg^2+^ that is added together with nucleotides caused strong effects on fusion independent of ISWI (Fig. S3g).

We conclude that ISWI, when stuck in individual nucleotide states, stiffens condensates. The HSS domain participates in this process, likely by forming bridges to a neighboring nucleosome. It is active hydrolysis, in other words the active cycling through all nucleotide states, that prevents ISWI from stably bridging nucleosomes.

### A theoretical framework to describe ISWI diffusion in condensed chromatin

We next derived a model for ISWI-chromatin interactions in condensates. The two DNA binding domains of ISWI, the ATPase and the HSS domain, are flexibly linked (Ludwigsen et al., 2013) and undergo large conformational changes during the ATPase cycle (Harrer et al., 2018; Leonard & Narlikar, 2015). These conformational changes might allow ISWI to make and break interactions with neighboring nucleosomes during the ATPase cycle.

When ISWI binds two nucleosomes on different DNA fibers at once, nucleosome condensates should stiffen, leading to slower condensate fusion. The drastically slower fusion velocities that we observed when ISWI was bound to ADP, AMPPNP or ADP-BeFx therefore indicate that ISWI can bridge nucleosomes in these nucleotide states (Fig. 3d, e). Of note, bridge formation is nucleotide-state dependent. Addition of nucleotide-free ISWI to condensates had only a modest impact on fusion velocity, suggesting that nucleotide-free ISWI only forms unstable bridges. Yet, it is still stably bound to chromatin as suggested by FRAP (Fig. S3a). Taken together, the results indicate that only one of ISWI’s domains interacts with chromatin in the nucleotide-free state.

ISWI FRAPs slowly in its AMPPNP, ADP-BeF_x_ and ADP-bound states, suggesting that ISWI interacts with nucleosomes with at least one of its binding domains in all these states. Intriguingly, ISWI FRAPs rapidly in the presence of ATP. It is thus the progression through ISWI’s ATPase cycle that repeatedly modulates the binding affinities of both binding domains. Hydrolysis-powered conformational changes would also repeatedly break up nucleosome-nucleosome bridges, explaining why addition of ATP prevents stiffening of condensates in condensate fusion experiments. We refer to this model as a “monkey bar” mechanism (Rudolph et al., 2018) as it results in an alternating release of one or the other binding site during the ATPase cycle (Fig. 4a).

**Figure 4.**
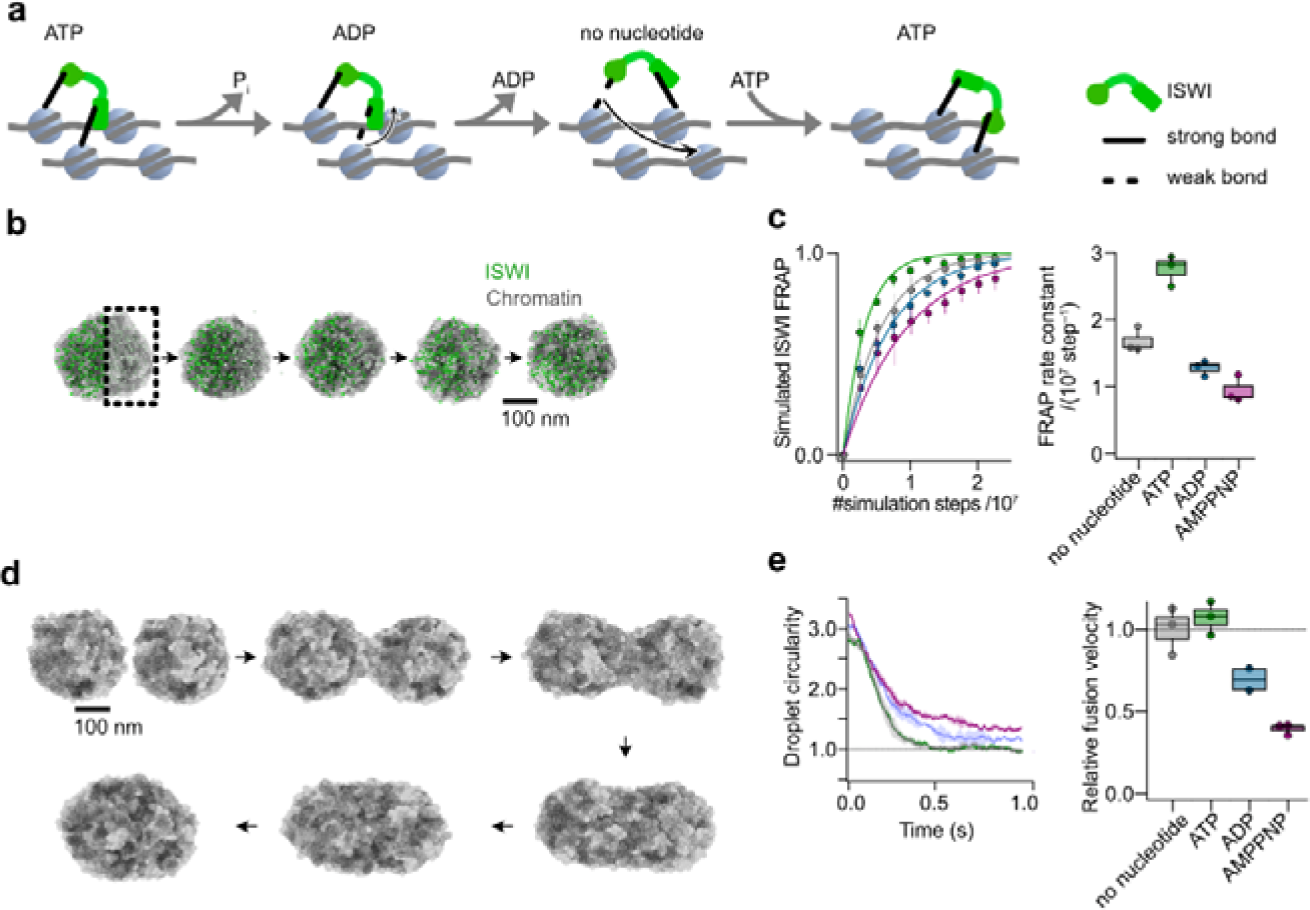
A simple model qualitatively explains experimental observations. **a,** ISWI possesses two nucleosome-interacting domains, which can bridge neighboring nucleosomes in dense chromatin. Via large nucleotide-induced conformational changes (not shown), the domains cycle through high and low affinity states towards nucleosomes, allowing ISWI to actively translocate through chromatin. **b,** Representation of a molecular dynamics simulation of ISWI FRAP in a chromatin condensate. ISWI particles on one side of an equilibrated condensate are switched to a bleached state at t=0 (square). Particle dynamics are then simulated to observe mixing of the two ISWI populations. **c,** ISWI FRAP was simulated as in b for 2.25*10^7^ timesteps in different nucleotide conditions with three replicates per condition. Left: Averaged traces of ISWI FRAP in different nucleotide conditions. Right: Rate constants of the simulated FRAP traces. **d,** Representation of simulated fusion experiments. Two equilibrated condensates were brought into proximity and allowed to fuse. For visual clarity only nucleosomes are shown here. **e,** Simulations of three fusion events for each nucleotide condition. Left: Averaged fusion traces with standard error. Right: Relative fusion velocities of the simulated traces.

ADP-bound and nucleotide-free ISWI display different behavior in fusion and FRAP experiments. ISWI in both states FRAPs slowly, whereas ISWI in the ADP but not the nucleotide-free state stiffens condensates. This difference suggests that the two interaction sites have individual nucleotide state-dependent binding affinities for nucleosomes. In both ADP-bound and nucleotide-free ISWI, one domain has a higher binding affinity than the other. The FRAP rate will be dominated by the higher affinity interaction. The fusion speed, on the other hand, also relies on the strength of the weaker affinity of the other domain. In the ADP-bound state, the weaker interaction domain therefore likely possesses an affinity that is higher than in the nucleotide-free state.

We formalized these findings in terms of a simple kinetic model (Fig. S4a). In the ATP state, ISWI stably binds with both domains to two nucleosomes. Upon hydrolysis, a conformational change weakens one of the interactions and leads to the release of one nucleosome. Subsequent ADP release switches the strengths of the two interactions, leading to rebinding of the first domain to a nucleosome, while releasing the second nucleosome. This alternating binding and unbinding of the two domains would enable ISWI to actively move through a dense mesh of nucleosomes and prevent ISWI from stably crosslinking neighboring fibers.

To test if such a simple, idealized model could recapitulate our observation that ATP hydrolysis allows fast FRAP and prevents condensate hardening in fusion experiments, we performed molecular dynamics simulations using Rouse chains to model chromatin arrays and describing all other interactions using Lennard-Jones potentials (Fig. S4b-d). Different nucleotide states of the remodeler were modeled by changing the depth of the respective interaction potentials.

In ISWI FRAP simulations (Fig. 4b, Movie S5), we observed enhanced recovery of the remodeler in the case of active ATP hydrolysis compared to remodeler that was stalled in any other nucleotide state (Fig. 4c). Like experiments, simulations of fusion (Fig. 4d, Movie S6) show no significant difference between ATP and nucleotide free conditions (Fig. 4e). The ADP and AMPPNP conditions are noticeably slower and do not reach full circularity within the simulated time. This again hints at the formation of ISWI-mediated stably bridged chromatin networks in the condensates that leads to a decrease in condensate fluidity.

## Discussion

One major result of our study is that the nucleosome remodeling enzyme ISWI can work largely unimpeded in folded and condensed nucleosome environments. This conclusion likely extends to other remodeling complexes given that their remodeling activities were minimally affected by histone hyperacetylation or trypsinization (Boyer et al., 2000; Logie et al., 1999). Our finding implies that all nucleosomes are similarly accessible for the enzyme in condensed arrays. As nucleosome sliding takes place in 1 bp increments (Deindl et al., 2013), each bp movement induces a rotation of the nucleosome by ∼36° relative to other nucleosomes. Sliding inside condensed arrays, therefore, necessitates that the chromatin structure is highly fluid and readily accommodates large, local structural changes imposed by nucleosome sliding.

ISWI critically depends on interactions to the H4 N-terminal tail and the nucleosome acidic patch for catalysis (Clapier et al., 2001; Dann et al., 2017; Gamarra et al., 2018). Incidentally, both these epitopes are also suggested to be important for array folding (Schalch et al., 2005). The fact that catalysis is not strongly impaired in condensed nucleosomes suggests that these epitopes are still dynamically available for ISWI.

Nucleosome concentrations in condensates reach ∼0.2 mM, a regime that nuclear enzymes also encounter *in vivo* (Hihara et al., 2012; Weidemann et al., 2003). We show that despite this high concentration, condensates remain viscoelastic materials and can accommodate dextrans with a Stokes radius equivalent to a 6 MDa protein complexes. Thus, even large remodeling complexes and other nuclear machines retain a chance to enter condensed chromatin, consistent with prior literature (Beaudouin et al., 2006; Erdel et al., 2015; Verschure et al., 2003; M. Zhang et al., 2022).

ISWI strongly enriches in chromatin condensates, further illustrating condensate plasticity. We anticipate that the same will be true for other chromatin factors simply because they possess high affinities for nucleosomes. It is as if chromatin is the scaffold in condensates to which client proteins like ISWI bind. This simple scaffold-client concept, often entertained in condensate biology (Ditlev et al., 2018), does not capture the complexity of the reaction, however. Inevitably, the client will affect chemical and physical properties of the scaffold. We found, for instance, that binding of ISWI hardens chromatin condensates, probably by crosslinking fibers. Remodelers, therefore, could serve structural purposes in addition to their well-known enzymatic activities (Korber & Becker, 2010).

The biophysical properties of nucleosome condensates varied with ambient conditions. Increasing Mg^2+^ concentrations, for instance, drastically stiffened condensates. As free Mg^2+^ increases during mitosis (Maeshima et al., 2018), stiffening of chromatin could facilitate sister chromatid condensation (Shimamoto *et al*., 2017). Moreover, changes in the extracellular milieu could rapidly elicit cellular responses through changes in viscoelastic properties of cellular components, including chromatin (Kroschwald et al., 2018; Munder et al., 2016).

Biomolecular condensates are suggested to affect enzyme activities through their unique chemical environments (Zhang et al., 2021). Performing enzyme kinetics in non-homogeneous solution is a challenge. Here, we have developed a FLIM-FRET nucleosome sliding assay to overcome such limitations. Its spatial resolution allows us to compare nucleosome sliding in- and outside of condensates. We predict that this kind of technology could become beneficial in the future for tracking enzymatic processes in cellular condensates in real time.

Enzymatic reactions inside biological condensates were suggested to benefit from high concentration of components (Zhang et al., 2021). However, rate enhancements would be expected for enzymes that have to work at subsaturating substrate concentrations. This is not the case for remodelers, which face nucleosome concentrations in the nucleus far above their *K*_D_, explaining why ISWI did not experience rate enhancements upon condensate formation.

Regardless of the mode and extent of chromatin phase separation occurring *in vivo* (Erdel et al., 2020; Erdel & Rippe, 2018; Keizer et al., 2022; Schneider et al., 2022), we view nucleosome condensates as useful model systems to study challenges enzymes face at the enormously high nucleosome concentrations in the nucleus. For instance, multivalent recognition of the nucleosome paired with the high nucleosome concentration could promote non-productive interactions (Mueller-Planitz et al., 2013). Given that nucleosome concentrations generally far exceed *K*_D_-values of chromatin enzymes, almost all enzyme molecules would be expected to be bound to chromatin at equilibrium. Our FRAP data support that notion, showing that ISWI becomes stuck in condensates if ATP cannot be turned over. This finding also helps explain why ATPase-dead variants of ISWI are easier to ChIP (Gelbart et al., 2005; Whitehouse et al., 2007) even though nucleotides only weakly modulate affinities (Leonard & Narlikar, 2015). How, then can these enzymes be mobile enough to rapidly diffuse through chromatin? ISWI could mobilize (“stir up”) chromatin by shuffling nucleosomes around, but we found no evidence in support of this possibility.

Instead, we propose the existence of a novel functionality of ISWI’s ATPase activity: besides nucleosome sliding, ATP hydrolysis may power conformational changes that pull the nucleosome binding domains of ISWI away from nucleosomes, leading to dissociation and subsequent re-association to other nucleosomes. Non-productive interactions could be dissolved in this fashion, and we speculate that some of the frequent pauses that ISWI remodelers take during sliding could be caused by unproductive binding (Blosser et al., 2009; Deindl et al., 2013). According to this mechanism, ISWI effectively remains bound to nucleosomes at any time, yet active ATP hydrolysis fuels its diffusion through chromatin. At the same time, hydrolysis prevents ISWI from crosslinking and thereby hardening condensates. This mechanism may be shared with other fly and yeast remodelers, which also became more dynamic in presence of ATP (Kim et al., 2021; Tilly et al., 2021). The mechanism however may not extend to Snf2H and Snf2L transfected into human cells, which were mobile before and after ATP depletion (Erdel et al., 2010). These remodelers may rely on a different mechanism, perhaps due to heteromerization (Oppikofer et al., 2017).

Our findings add another facet to why loss of function mutations can induce dominant negative phenotypes or be cancer-associated (Clapier et al., 2020). Half of SMARCA4 mutations in cancers lie in the ATP cleft and these mutants show slower FRAP (Hodges et al., 2018). Expression of ATPase dead BRM in *Drosophila* caused peripheral nervous system defects, homeotic transformations, and decreased viability (Elfring et al., 1998). All phenotypes were dominant negative (Elfring et al., 1998; Hodges et al., 2018) and might be caused in part by changes in chromatin dynamics, and not exclusively by disruption of canonical remodeler functions. Changed dynamics of biomolecular condensates may in turn disturb biological functions (Li et al., 2020; Shi et al., 2021). It is our hope that our tools and concepts will help open new avenues for mechanistic dissections and development of therapeutics.

## Methods

### Cloning

#### Cloning of GST-sfGFP and 6xHis-TEV-ISWI-3C-STREP-sfGFP

Full length super fold GFP (Pédelacq et al., 2006) sequence was amplified from the plasmid generously provided from Ökten group at TU Munich (pFMP248). The PCR product and pET41b(+) were digested with *BgIII* and *EcoRI*, ligated and the resulting plasmid containing GST-sfGFP construct (pFMP241) amplified in *E. coli DH5α*. Correct sequence was confirmed by Sanger sequencing.

To make 6xHis-TEV-ISWI-3C-STREP-sfGFP construct, the PCR amplified sfGFP sequence was first subcloned into vector pUC57 containing synthesized (GenScript) C-terminus for 3C-STREP-KKCKK-GFP11 (pFMP249). For that purpose, both insert and vector were digested with *MslI* and *HindIII*. *MslI* cuts inside of GFP11 (extreme C-terminal sequence of sfGFP) and leaves six residues. The obtained plasmid was then digested with *XbaI* and *HindIII* and ligated into pFMP210 digested with the same enzymes, and pFMP244 was produced and amplified in *E. coli DH5α*. Correct sequence was confirmed by Sanger sequencing.

### Protein expression and purification

#### GFP expression and purification

Plasmid containing GST-GFP (pFMP248) was transformed into *E.coli* BL21 (DE3) and incubated on selective plate at 37 °C overnight. A single colony was inoculated into 15 mL LB media supplemented with ampicillin and grown overnight at 37 °C. The next morning the culture was diluted to OD=0.05, grown at 37 °C until OD=0.6 and induced with 0.2 mM IPTG. Protein was expressed overnight at 20 °C with shaking. The bacterial cultures were harvested (rotor SS34, 14000 rpm, 10 min) and pellet was stored at -80 °C. Pellet from 485 mL expression was resuspended in 20 mL of GST binding buffer (25 mM Tris-HCl pH=7.5, 150 mM NaCl, 1 mM EDTA, 0.5 mM DTT, 20 mM imidazole, 1 mM PMSF, 30 µL of each leupeptin, pepstatin, aprotinin and one tablet complete EDTA-free protease inhibitor cocktail (Roche)). Cells were disrupted in French press (3× 1500 PSI) and sonication (12×10 s on, 20 s off, amplitude 22%). The suspension was centrifuged (SS34 rotor, 19000 rpm, 30 min, 4 °C) and to the supernatant 1 mL of prewashed GST-beads was added (50% slurry). The suspension was incubated for 90 min at 4 °C with gentle mixing. The beads were removed with centrifugation (1000 rpm, 1 min, 4 °C) and washed five times with 10 mL binding buffer. Finally, beads were resuspended in 1.5 mL elution buffer (50 mM Tris-HCl pH=8.0, 10 mM reduced glutathione, 5% glycerol) and incubated on rotating wheel for 30 min at 4 °C. The beads were removed by centrifugation and supernatant aliquoted, frozen in liquid nitrogen and stored at -80 °C.

#### Expression and purification of ISWI, ISWI-GFP, ISWI_ΔHSS_ and ISWI E257Q

Corresponding plasmids containing *D. melanogaster* ISWI (pFMP210 for 6xHis-TEV-ISWI, pFMP244 for 6xHis-TEV-ISWI-3C-TREP-sfGFP, pFMP114 for 6xHis-TEV-ISWI26−648 (*ISWI_ΔHSS_*), pFMP110 for 6xHis-TEV-ISWI E257Q) were freshly transformed into BL21 Star *E. coli* and plates were incubated at 37 °C overnight. The bacterial lawn from one plate was harvested and used to inoculate two l of LB media. Cultures were grown (37 °C, 130 rpm) until OD 0.5-0.6 when they were induced with 1 mM IPTG. The expression went overnight at 18 °C with shaking. The bacterial cultures were harvested (6000×g, 10 min, 4 °C). Pellets were gently rinsed with cold water and stored at -80 °C. Except for ISWI_ΔHSS_, whose purification is described in previous publications (Harrer et al., 2018; Mueller-Planitz et al., 2013), all ISWI proteins were purified as follows. Cells were thawed in a palm of a hand with occasional vortexing and resuspended in HisA buffer (50 mM Tris-HCl pH=7,4, 300 mM NaCl, 0.5 mM DTT) supplemented with 20 mM imidazole (pH=7.4), 1 tablet complete-EDTA free protease inhibitors (Roche), as well as leupeptin (1 mg/L), pepstatin (0.7 mg/L) and aprotinin (1 mg/L). Benzonase (Merck Millipore 1016540001 100000; 5 μl per liter of culture) and lysozyme (tip of a spatula) were added and suspension was sonicated on ice (Branson sonifier, 6×10 seconds on, 10 seconds off, amplitude 25%). The homogenized bacteria were then cracked with six runs on Microfluidizer LM10 at 1200 bar. Lysates were centrifuged (19000 rpm, JA25.50 rotor, 4 °C, 30 min), supernatants filtered through 0.45 µm syringe filter and loaded onto His Trap (5 mL, GE Healthcare) equilibrated with 5% HisB buffer (100% HisB; 50 mM Tris-HCl pH=7,4, 300 mM NaCl, 400 mM imidazole). The column was washed with 10 column volumes (CV) 5% HisB, then with 6 CV 10% HisB and 1 CV 20% HisB. ISWI was then eluted with 20-100% HisB gradient over 10 CV. ISWI containing fractions were pooled, TEV protease added (0.1 mg for every 8 mg of ISWI) and the mixture was dialyzed (Spectra/Por dialysis membrane, cutoff 12000-14000 kDa) overnight against dialysis buffer (15 mM Tris-HCl pH = 7.4, 150 mM NaCl, 1 mM DTT). To remove His-tagged TEV protease, cleaved-off 6xHis-tag and uncleaved 6xHis-TEV-ISWI, a second nickel-affinity chromatography was performed (His Trap, 5 mL, GE Healthcare; equilibrated in 10% HisB buffer). Sample was loaded and unbound protein further washed out with 10% HisB. The flowthrough was pooled and its conductivity reduced by slow dilution with two volumes of 2% MonoS B buffer (100% MonoS B =15 mM Tris-HCl pH=7.4, 2 M NaCl, 1 mM DTT), which was diluted with MonoS A (15 mM Tris-HCl, 1 mM DTT) and filtered through 0.2 µm syringe filter. A MonoS column (5/50 GL GE Healthcare 1 mL) was equilibrated with 2% MonoS B. After loading, the protein was eluted with 2-30% MonoS B (10 CV) and 30-100% MonoS B (4 CV). Fractions containing ISWI were pooled together, concentrated and loaded onto the HiLoad Superdex200 (GE Healthcare, 120 mL) exclusion column equilibrated with GF buffer (50 mM Hepes KOH pH=7,6, 0.2 mM EDTA pH=8, 200 mM KOAc, 10 mM DTT). ISWI-containing fractions were pooled and protein was concentrated to ∼5 mg/mL. Molar concentration was determined by using calculated extinction coefficient (Expasy Protparam tool). Proteins were aliquoted, frozen in liquid nitrogen and stored at -80 °C.

#### Expression and purification of histones

Codon optimized *D. melanogaster* histones were expressed and purified as previously described (Klinker, Haas, et al., 2014). Corresponding plasmids (pFMP128 for H2A, pFMP129 for H2B, pFMP186 for H3, pFMP187 for H4, pFMP269 for H2AK119C, pFMP270 for H3C111A, pFMP268 for H4T1C) were freshly transformed into BL21 Star *E. coli* and incubated at 37 °C overnight. Bacterial lawns from one plate were then used to inoculate two liters of LB media. Cultures were grown (37 °C, 130 rpm) until OD reached 0.5-0.6 when they were induced with 1 mM IPTG. The expression went for three hours at 37 °C with shaking. The bacterial cultures were spun down (6000xg, 10 min, 4 °C). Pellets were gently resuspended with cold water, transferred to a 15 ml Falcon tube. Samples were centrifuged and the pellets stored at -80 °C. Cells were thawed in a palm of a hand with occasional vortexing and resuspended in SA buffer (40 mM NaOAc pH=5.2, 1 mM EDTA pH=8, 10 mM lysine) supplemented with 6 M urea, 200 mM NaCl, 1 mg/mL aprotinin, 1 mg/mL leupeptin, 1 mg/mL pepstatin, 1 mM PMSF and 5 mM β-mercaptoethanol. Benzonase (Merck Millipore 1016540001 100000; 5 μl per liter of culture) and lysozyme (tip of a spatula) were added and suspension was sonicated on ice (Branson sonifier, 15 seconds on, 30 seconds off, amplitude 30%, effective sonication time 20 minutes). The homogenized bacteria were then cracked with six runs on Microfluidizer LM10 with 1200 bar. Lysates were centrifuged (19000 rpm, JA25.50 rotor, 4 °C, 30 min), supernatant filtered through 0.45 µm syringe filter and loaded onto a HiTrap Q HP column (5 ml, GE Healthcare) that was stacked on top of a SP column (5 ml, GE Healthcare) equilibrated with 20% buffer B (buffer A: 40 mM NaOAc pH=5.2, 1 mM EDTA pH=8, 10 mM lysine, 7.5 M urea, 5 mM DTT; buffer B: 40 mM NaOAc pH=5.2, 1 mM EDTA pH=8, 10 mM lysine, 7.5 M urea, 5 mM DTT, 1000 mM NaCl). Samples were applied to the stacked columns. The columns were washed with 20% buffer B (1 CV). The Q column was removed and SP column further washed with 25% buffer B (3 CV) and 30% buffer B (3 CV). Histone was eluted with 30-40% buffer B gradient (5 CV), 40-80% buffer B gradient (7 CV), 100% buffer B (3 CV). Pooled fractions were dialyzed (SpectraPor MWCO 3,500 kDa) three times against 5 L of miliQ water. Purity was analyzed on SDS-PAGE and concentration determined from A280 absorption. Histones were aliquoted (1 mg per aliquot), flash frozen in liquid nitrogen and stored at -80 °C or -70 °C. Histones were lyophilized before use. Lyophilized histones were stored at -20 °C.

#### Assembly and purification of octamers

Lyophilized histone aliquots were dissolved in an unfolding buffer (20 mM Tris-HCl, 7 mM guanidinium-HCl, 20 mM DTT) to 4 mg/ml for 10 minutes in a thermoblock (24 °C, 600 rpm). Solutions were spun down in table top centrifuge (10 minutes, full speed, 4 °C), and supernatants transferred to fresh tubes and kept on ice until dialysis. Histone concentrations were remeasured in unfolding buffer by measuring OD280 and corrected for purity as assessed from SDS-PAGE gel. Histones were mixed in molar ratio H2A:H2B:H3:H4=1.4:1.4:1:1. The histone mixture was transferred into dialysis membranes (Roth E658.1 MWCO: 4000-6000) which were soaked in water for one hour and rinsed with refolding buffer (10 mM Tris-HCl pH=7.5, 2 M NaCl, 1 mM EDTA, 5 mM β-mercaptoethanol). The mixture was then dialyzed three times against 1 L of refolding buffer, with the second dialysis step being overnight. Lastly, octamers were purified by size exclusion chromatography (HiLoad 16/60 Superdex 200 prep grade), concentrated to 4 mg/mL (Amicon 15 mL 30 kDa), aliquoted, flash frozen in liquid nitrogen and stored at -80 °C or -70 °C.

#### Histone labeling (H2AK119C-atto565, H4T1C-cy3)

Lyophilized histone aliquots were dissolved in a labeling buffer (7 M guanidinium-HCl, 20 mM Tris-HCl pH=7.5, 5 mM EDTA, 0.7 mM TCEP) to final concentration 0.2 mM and incubated two hours to reduce all cysteines. Cyanin-3-maleimide (Lumiprobe) was dissolved in DMSO to final concentration 100 mM and added to solution in 5.7 fold excess. Atto-565 maleimide (Atto-tec) was dissolved in DMF to final concentration 100 mM and added to solution in 13.2 fold excess. Histones were incubated with the dye for 14 hours (3 hours for cy3 labeling) on rotating wheel at room temperature covered with aluminum folium. To stop the labeling reaction, β-mercaptoethanol was added to final concentration to 340 mM. For cy3 labeling, reaction was stopped by adding DTT to final concentration 20 mM. Unreacted dye was partially removed by several rounds of ultrafiltration (15 mL 10K Amicon, Millipore) and successive dilutions with labeling buffer (this step was omitted for Cy3 labeling). The labeling efficiency was assessed with SDS-PAGE and fluorescence imaging.

### Synthesis of DNA for nucleosome arrays

To distinguish remodeling of individual nucleosomes in a nucleosome array, we previously cloned a DNA with unique restriction sites in 13 of 25 Widom-601 positioning sequences with a 197 bp repeat length (pFMP233) (Ludwigsen et al., 2018) (Fig. S1c). Here we cloned a second variant of this DNA by placing the 13 unique restriction sites on the other end of the canonical 601 sites, essentially inverting the unique restriction sites with regards to their positioning inside of the array, yielding pFMP232 (Fig. 1a). To this end, thirteen 601 repeats were synthesized (GenScript) in which naturally occurring AvaI and AluI sites in the canonical 601 sequenced were replaced with unique restriction sites. A unique AvaI restriction site was added to the 3’-end (pFMP226). The 13-mer was extended to a 25-mer as follows. pFMP226 was completely digested with XbaI, EcoRI and AseI in buffer II (NEB) and the insert was subcloned into pUC18, which was fully digested with XbaI and EcoRI, yielding pFMP236. A plasmid containing 25×197 bp canonical Widom-601 repeats (pFMP166) was subjected to partial AvaI digestion. The band corresponding to a 12mer repeat (2354 bp) was gel purified and cloned via ligation into AvaI digested pFMP236, yielding pFMP232 (Fig. 1a).

### Preparation of DNA for nucleosome arrays

25mer and 13mer nucleosome arrays were prepared as published previously (Ludwigsen et al., 2018). Briefly, plasmids carrying 25 (pFMP232, pFMP233) or 13 (pFMP226) consecutive copies of 197 bp with modified 601 sequence were transformed into DH5α *Escherichia coli* strain and purified from four liters of culture by using plasmid DNA purification kit Nucleobond® PC 10000 (Macherey-Nagel). Three mg of the plasmids were digested with EcoRI HF (0.25 U/µg DNA) and HincII (0.6 U/µg DNA) in CutSmart Buffer at 37°C for 3 h to cut out the arrays from the plasmids. When the digest was complete, restriction enzymes were heat-inactivated by incubation at 65°C for 20 min. Tubes were put on ice before AseI was added (0.5 U/ µg DNA) and then incubated at 37 °C for four h. After digest was complete, DNA was purified via phenol/chloroform extraction and ethanol precipitated in presence of NaOAc. Finally, it was resuspended in TE buffer (10 mM Tris-HCl pH=8, 1 mM EDTA pH=8) and stored at -20 °C.

### Preparation of DNA for mononucleosomes

DNA fragments containing the 601 sequences were amplified by large-scale PCR (Yang & Narlikar, 2007). Primers were obtained from Sigma. For primer sequences, see Table S1. The PCR was cleaned up by precipitating the plasmid template by addition of ½ of volume 30% PEG 8000 (w/w) in 30 mM MgCl2. The PCR product was then precipitated by adding the same volume of propan-2-ol. The pellet was washed with 70% (v/v) cold ethanol, resuspended in TE buffer and stored at -20 °C.

### Chromatin assembly and purification

#### Assembly of 25- and 13mer arrays

The optimal molar octamer:601 ratio was identified by performing small scale test assemblies, where purified octamers were titrated to 7.5 µg digested plasmid (Ludwigsen et al., 2018). Preparative assembly contained 100-500 µg digested plasmid (100 ng/μL of 601-array DNA, which corresponds to 150 or 200 ng/μL of total digested plasmid containing 25mer or 13mer, respectively). It also contained corresponding amounts of purified octamers, 2 M NaCl, 10 mM Tris-HCl pH=7.6, 1 mM EDTA pH=8, 1 mM DTT. Reactions were transferred into dialysis membrane and underwent salt gradient dialysis: 3 L of low salt buffer (50 mM NaCl, 10 mM Tris-HCl pH=7.6, 1 mM EDTA pH=8, 1 mM DTT) was pumped into 1 L of high salt buffer (2 M NaCl, 10 mM Tris-HCl pH=7.6, 1 mM EDTA pH=8, 1 mM DTT) containing the dialysis bag over a period of 24 hours. To maintain the constant volume, buffer was simultaneously pumped out with the same speed. Assemblies were then dialyzed against 1 L of low salt buffer before they were precipitated by addition of equal volume of precipitation buffer (10 mM Tris–HCl pH 7.6, 7 mM or 10 mM MgCl2 for 25mer or 13mer, respectively). Pellets were resuspended in TE buffer (10 mM Tris-HCl pH=7.6, 1 mM EDTA pH=8) and quality controls of chromatin array were performed as described (Ludwigsen et al., 2018) (Fig. S1a). Chromatin concentrations were approximated by UV assuming that 1 OD at 260 nm equals 50 ng/µL of DNA.

#### Mononucleosomes

The optimal molar octamer:DNA ratio was first identified by titrating octamers to DNA. Preparative assemblies contained DNA (200 ng/µL), purified octamers, 2 M KCl, 20 mM Tris-HCl pH=7.7, 10 mM DTT. Reactions were transferred into Slide-A-lyzer 7k Mini and underwent salt gradient dialysis in 200 mL of Mono2000 buffer (2 M KCl, 20 mM Tris-HCl pH=7.7, 0.1 mM EDTA pH=8, 1 mM DTT) to which 1 L of Mono0 buffer was pumped (20 mM Tris-HCl pH=7.7, 0.1 mM EDTA pH=8, 1 mM DTT) over 24 hours. To maintain constant volume, buffer was simultaneously pumped out with the same speed. Assemblies were dialysed against Mono0 buffer and purified over 10-30% (w/w) glycerol gradient. Concentrations were determined as above for arrays.

### Quality controls of assembled chromatin

#### Agarose gel of nucleosome arrays

200 ng of before and after Mg-precipitation was analyzed by agarose gel electrophoresis (0.7%).

#### NotI digestion of nucleosome array

200 ng of arrays were digested in EX50 buffer (10 mM Hepes–KOH pH 7.6, 50 mM KCl, 1.5 mM MgCl2, 0.5 mM EGTA) with Not1 (20 U/μL) in total volume of 15 μL for three hours at 26 °C. Digestion was analyzed on 1.1% agarose.

#### BsiWI digestion of nucleosome array

250 ng of an arrays were digested in buffer (25 mM Hepes–KOH pH 7.6, 0.1 mM EDTA, 50 mM NaCl, 10% glycerol, 2 mM MgCl2) with BsiWI (10 U/μL) for 1 hour at 26 °C in total volume of 20 µL. The digestion was stopped with addition of SDS (final concentration 0.4%) and EDTA (final concentration 20 mM) followed by Proteinase K (1 mg/mL) treatment in total volume of 30 µL for three hours at 65 °C or overnight at 37 °C. DNA was ethanol precipitated and analyzed on 1% gel.

### Analytical ultracentrifugation

Sedimentation velocity (SV) experiments of purified, reconstituted arrays were conducted at 20 °C in a Beckman Coulter Optima XL-I analytical ultracentrifuge (Palo Alto, CA) using an An-50 Ti rotor. Samples contained 21.6 ng/µL (6.9 nM) of 25mer and were dissolved in buffer (1 mM Tris-HCl pH 8.0, 0.01 mM EDTA pH 8.0, 0.01 mM DTT, 50 mM NaCl, 0.2 mM or 1.7 mM MgCl2). Samples (360 µl) were loaded into 12 mm charcoal-filled epon double sector centerpieces. A rotor speed of 22,000 rpm was selected and absorbance optics scans at a wavelength of 258 nm were collected every second until sedimentation was complete. Data were analyzed using the c(s) model in SEDFIT which directly models the sedimentation boundary as a continuous distribution of discrete, non-interacting species (Schuck, 2000). Buffer density and viscosity as well as sample partial specific volumes were calculated using UltraScan III (Demeler & Gorbet, 2016).

### Negative stain electron microscopy

Quantifoil R2/1 Cu200 C2 grids were plasma cleaned for 20 s at 20 mA (GloCube, Quorum). 3.5 µL of sample containing 1-4 ng/µL (0.3-1.3 nM) of 25mer in buffer (3 mM Tris-HCl pH 8.0, 0.03 mM EDTA pH 8.0, 0.03 mM DTT, 50 mM NaCl, 0.2/1.7 MgCl_2_) was applied, incubated 30 s, then hand blotted. Grids were negative stained with 2× 3.5 µL of 2% uranyl acetate, and hand blotted after 30 s for each stain application. Images were collected using an FEI Morgagni 100 keV TEM with a SIS Megaview III 1k CCD, at a nominal magnification of 56,000×.

#### Analysis of electron micrographs

Outlines of single particles were determined with a trainable Weka segmentation in ImageJ (Fiji). Feret’s diameter (the maximum distance between two parallel tangential lines), and circularity were calculated for this outline using equation 1.

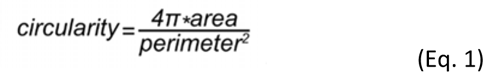

### Restriction based nucleosome sliding assay

Remodeling assays contained 4 nM 25mer arrays (100 nM total nucleosome concentration), 200 nM ISWI, ATP regeneration system (6 mM phosphoenolpyruvate, 15.5 U/mL pyruvate kinase/lactate dehydrogenase, 1 mM DTT), indicated magnesium concentration and 5 µM Mg-ATP in remodeling buffer RB (25 mM Hepes pH=7.6, 0.1 mM EDTA, 50 mM NaCl, 10% (v/v) glycerol, 0.2 mg/mL BSA). Beforehand, chromatin arrays were dialyzed overnight into 10 mM Tris pH=7.6 at 4 °C. The reaction was started with addition of ISWI and incubated at 26 °C. At different time points, 20 µL aliquots were taken and remodeling was quenched with apyrase (50 mU, 1 min, 26 °C). The Mg^2+^ concentration was supplemented to a final concentration of 1.7 mM MgCl_2_ for all samples. Samples were digested with 10 U/125 ng array of BamHI for 90 min at 26 °C. The digestion was stopped with addition of 20 mM EDTA and 0.5% SDS, followed by Proteinase K (final concentration 0.5 mg/mL) treatment for three hours at 37 °C. Samples were ethanol precipitated and separated on an agarose gel (0.7% to 0.9% in 0.5xTBE, 20 cm). Gels and running buffers contained 0.5 µg/mL EtBr. The bands were analyzed with *AIDA Image Analyzer Software* and the percentage of *Cut*-DNA calculated. The dependence of percentage of Cut-DNA versus time was fitted in R into to a single exponential function (Eq.2):

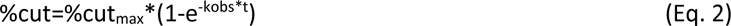

### ATPase assay

A NADH-oxidation coupled ATPase assay was performed as described (Mueller-Planitz et al., 2013). Briefly, 30 µl reactions were assembled on a 384-well plate (Greiner, 781101), containing 100 nM ISWI and 0.1/0.5/1.33 µM mononucleosomes (0N60) in remodeling buffer supplemented with ATP-regeneration system and 0.6 mM NADH. Each sample was measured in technical triplicates. Reactions were started by addition of 1 mM Mg-ATP and NADH absorbance was monitored at 340 nm in a plate reader (Biotek PowerWave HT) at 26°C. Absorbance readings between ten and 20 minutes were fit to a linear function.

### Phase separation of nucleosome arrays and imaging of formed chromatin condensates

All phase separation experiments were performed in phase separation buffer (PSB) (25 mM Hepes-KOH pH 7.6, 0.1 mM EDTA, 50 mM NaCl, 10% glycerol, 1 mM DTT, 0.2 g/L BSA) supplemented with 1 mM MgCl_2_ (PSB1) or 5 mM MgCl_2_ (PSB5). Phase separation was induced by mixing equal volumes of nucleosome arrays diluted in TE buffer with 2× PSB. The formed chromatin condensates (4-10 µL of sample) were incubated at room temperature for a minute and then loaded onto the imaging chamber made out of double-sided tape. Double-sided tape was pierced with a hole puncher before and sample was deposited into the hole. Cover slips were pretreated with 20 µL of BSA solution (25 mM Hepes-KOH pH 7.6, 0.1 mM EDTA, 50 mM NaCl, 10% glycerol, 1 mM DTT, 100 mg/mL BSA) for one minute. The imaging chamber was sealed with nail polish, and unless stated otherwise, spun onto the cover slip (1 min, 1000g, room temperature). Samples we then imaged on a widefield microscope (Zeiss Axiovert).

#### Analysis of phase diagram

Condensates were detected by Trainable Weka segmentation in ImageJ, their surface was calculated as a percentage of a total field of view surface and plotted in form of a heatmap in R.

### Confocal imaging

Confocal and FLIM images were performed with a TCS SP8 X FALCON confocal head (Leica Microsystems, Wetzlar, Germany) mounted on an inverted microscope (DMi8; Leica Microsystems). For confocal imaging, a white light laser was used as excitation source (561 nm or 633 nm as necessary). Single photons were collected through a 40×/1.3 NA oil-immersion objective and detected on Hybrid Detectors (HyD) (Leica Microsystems) with a 570 – 610 nm, and 650 – 707 nm spectral detection window as necessary. Sequential excitation was performed to avoid potential crosstalk between the fluorophores.

### FITC-dextran partitioning in chromatin condensates

Condensates were formed in a solution containing 100 nM 13mer (final concentration 90 nM), PSB2 supplemented with 2.5 mM DTT and incubated for five minutes at room temperature when the FITC-dextrans (Sigma) diluted in water were added to final concentration of 0.1 mg/mL. Samples were prepared as described above and imaged on Leica laser scanning confocal microscope after 1 h incubation at room temperature.

Images were analyzed in ImageJ, where the dextran partitioning coefficient was determined for individual condensates as a ratio of fluorescence inside the condensate and background fluorescence. Molecular weights of dextrans were converted into Stokes radii with an online tool (https://www.fluidic.com/toolkit/hydrodynamic-radius-converter/).

### ISWI colocalization experiment

Chromatin and ISWI colocalization experiment was performed with 40 nM of unlabeled 25mer, 10 nM of 25mer-Cy3 and 1.125 µM ISWI-GFP/GFP-GST in PSB5. Condensates were induced by adding Mg^2+^ (5 mM) either after or before addition of GFP-GST or ISWI-GFP.

A standard curve for mean Gray value dependence on ISWI-GFP concentration was obtained from ISWI-GFP dilutions in PSB5. Different microscope settings were used to image lower and higher dilutions. ISWI-GFP concentration was then determined inside condensates and in a surrounding solution for 90 nM 13mer, 234 nM ISWI-GFP, 1 mM Mg-ATP, ATP-regeneration system, PSB1 and for 45 nM 25mer, 125 nM 0N60 mononucleosomes, 625 nM ISWI-GFP, 1 mM Mg-ATP, ATP-regeneration system, PSB5.

### Restriction enzyme accessibility nucleosome sliding assay adapted to chromatin condensates

A *KpnI* site was used to compare nucleosome sliding in 25mer arrays fully dissolved or after condensate separation. A remodeling assay contained 15 nM 25mer, 750 nM ISWI, ATP regeneration system and 1 mM Mg-ATP in PSB0.2/5, in total reaction volume of 20 µL. Reaction was started with 2 µL of ISWI (Fig. 2d) or ATP (Fig. S2c) and incubated at 26 °C. Before the reaction was started, 6 µL of the reaction mixture were checked under the microscope for chromatin condensates. At different time points, 1 µL of reaction was quenched with 45 µL of quenching solution (10 mU/µL apyrase in apyrase reactions buffer, containing 1.8 µL 2 mM MgCl_2_ for low magnesium reactions), incubated at 26 °C for 15 min and after that kept on ice. After all time points were quenched, the reaction mixture was again checked under the microscope for the presence of condensates. To each tube, 2.5 µL of *KpnI* was added and incubated at 26 °C for 30 min. Cleavage was detected as above for the restriction-based nucleosome sliding assay, except that ImageJ was used for quantification.

### FLIM-FRET

#### Slide preparation

For end point assays (Fig. 2f), 45 nM 25mer (1125 nM nucleosome concentration), 125 nM labeled mononucleosomes, 625 nM ISWI, ATP regeneration system and 1 mM Mg-ATP were mixed in PSB5 in total volume of 10 µL. Samples were incubated at room temperature for 15 minutes, 6 µL was transferred on a slide (see above for slide preparation), spun as above, and imaged after four hours. For time lapse assays (Fig. 2g), 40 µL of the reaction mixture was loaded into the channel of imaging chamber (Ibidi μ-Slide VI^0.5^ Glass Bottom 80607). The imaging chamber was spun down and mounted on the microscope. Lifetimes were measured for two min before addition nucleotides. Then, 120 µL of 1 mM Mg-ATP or 1 mM Mg-AMPPNP solution, dissolved in identical buffer and supplemented with ATP regeneration system, was filled into one of the reservoirs without removing the chamber from the microscope or stopping the imaging. Nucleotide solution then replaced solution above the condensates by gravity flow. Time lapses were recorded for two h. Finally, time lapses in Fig. 2h and Fig. S2h contained 1125 nM unlabeled mononucleosomes or 45 nM 25mer, 125 nM FRET mononucleosomes and ATP regeneration system in PSB5 in total volume of 18 µL. Two µL of 10× Mg-ATP solution were added to a final concentration 1 or 5 mM, the mixture was loaded into a channel of the imaging chamber and imaged two to four min after ATP addition.

#### Image acquisition

For FLIM, the same system described in “Confocal Imaging” was used. The white light laser delivered 80 MHz repetition rate at 561 nm. Arrival time of single photons was measured with the included FALCON module. The FLIM acceptor photobleaching image kept the same parameters as the confocal one with 12 frames accumulations instead. The other FLIM images and movies size was set to 256 × 256 pixels. A 3-fold zoom factor was applied, giving a pixel size of 0.380 μm and an image size of 97 × 97 μm. Pixels number was decreased to favor imaging speed in the time-lapses. Because the statistical determination of the distribution of single photon arrival times requires a minimum number of photons, 60 frames were acquired at 2.34 Hz for each TCSPC recording, for a total time of around 26 s. Corresponding to a scanning speed of 600 Hz. Time-lapses were recorded for at least 15 min with a time point every 2 min.

#### Analysis

FLIM image analyses were performed in the LAS X software and with a home-made MatLab code (available on request). Lifetime calculations were based on the Phasor approach (Digman et al., 2008). Phase and modulation lifetimes were calculated using the Fourier sine and cosine transforms of the lifetime images. The FRET efficiency (E_FRET_) was calculated according to Eq. 3:

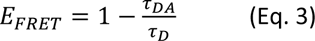

where τ_DA_ is the lifetime of the donor-acceptor sample, and τ_D_ is the lifetime of the donor alone. Results were expressed as mean ± SD. Lifetimes images shown were done using the phasor approach to maximize the number of photons while keeping a good image resolution. For the time lapse, the linear part of lifetime changes was fitted to a linear function to obtain initial velocity.

### Holotomography

Slide with 25mer condensates containing ISWI in PSB5 buffer was prepared as described above. Refractive index images were collected on 3D Cell Explorer-fluo (Nanolive) equipped with dry objective (60× magnification, 0.8 numerical aperture) and low power laser (λ = 520 nm, sample exposure 0.2 mW/mm2). 96 slices were collected for a field depth of 30 µm. Software Steve v.1.6.3496 (Nanolive) was used to collect, view and export images. Data were exported as tiff files (floating values of RI) and further processed in ImageJ (Fiji). Eq. 4 was used to calculate chromatin mass concentration in condensates, where n = refrective index, c = mass concentration and dn/dc is a refractive index increment. We assumed c_condensate_ >> c_solution_ and dn/dc to be 0.185 mL/g.

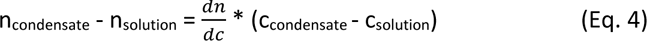

### ISWI-GFP and 25mer-Cy3 FRAP

Four µL of 2.5× mixture of unlabeled chromatin, labeled chromatin and ISWI-GFP were mixed with six µL 1.7xPSB5 containing 1.7xnucleotide. Final experimental conditions were: 76 nM 25mer, 1.75 nM Cy3-25mer, 1 µM ISWI-GFP, 0.77 mM Mg-nucleotide, PSB5, ATP regeneration system (Fig. 3b); 15 nM 25mer-Cy3, 375 nM ISWI-GFP, PSB5, 1 mM Mg-ATP/no nucleotide (Fig. S3a); 100 nM 25mer, 4.5 nM 25mer-Cy3, 1.3 µM ISWI-GFP, PSB5, ATP regeneration system, 1 mM Mg-ATP (Fig. S3b). The sample was incubated for 1 min and then loaded onto the slide, spun, sealed as above and imaged within 30 minutes of preparation.

Images were acquired at 26°C with a 63X glycerol immersion objective on a Leica Sp5 confocal microscope equipped with Argon 488 nm and DPSS 561 nm lasers. For ISWI-GFP FRAP, 10 frames (512×512 pixel) at 1.2 s intervals were taken as a pre-bleach reference, followed by a single 1.2 s bleaching pulse targeted to a circular region within ≥ 3 droplets at once. After bleaching, 89 frames were taken at 1.2 s intervals to measure fluorescence recovery. For H4-Cy3 FRAP, 10 frames (512×512 pixel) at 1.2 s intervals were taken as pre-bleach reference, followed by four 1.2 s bleaching pulses targeted to a circular region within ≥ 3 droplets at once. After bleaching, 20 frames were taken at 30 s intervals to measure fluorescence recovery. For both cases, brightfield images were also collected in parallel.

All images were processed using Fiji (Schindelin et al., 2012). First, drift was corrected using MultiStackReg package (https://biii.eu/multistackreg) by calculating transformation matrices from brightfield images (code available on request). Bleach, control (within droplet but outside the bleached region) and background Region Of Interest (ROIs) were manually defined, and average fluorescence intensity was measured. Intensities were normalized using the easyFRAP web tool (Koulouras et al., 2018) to generate FRAP curves with full scale normalization. Normalized FRAP curves from different droplets within the same experiment were considered as technical replicates. FRAP curves from different experiments were averaged and reported together with standard error of the mean (SEM). Plots were generated using R – version 4.2.1 (https://www.R-project.org/) (R Core Team, 2022).

### Controlled condensate fusion with optical tweezers

Four µL of chromatin ISWI mixture (225 nM 13mer, 585 nM ISWI, volume was made up with TE pH=7.6) was gently mixed with 6 µL 1.7xPSB1 containing 1.7xnucleotide. Final experimental conditions were: 90 nM 13mer, 234 nM ISWI, 1 mM Mg-nucleotide and ATP regeneration system in 1×PSB1. Data on fusion velocity with different ISWI concentrations (Fig. S3e) were collected in 1×PSB1 supplemented with 5 mM DTT. Of note, with an ATP-regeneration system present, slow fusion was detected with AMPPNP and ADP-BeFx, presumably due to ADP contamination present in nucleotide preparations. Samples were incubated for 1 min and then loaded onto prepared slides. Optical tweezer experiments were carried out on a dual-trap C-Trap (Lumicks, Amsterdam). For controlled fusion of condensates, a single condensate was trapped in each of the optical traps at minimal laser power, resulting in trap stiffnesses on the order of ∼0.001 pN/nm. The traps were approached in 20 nm steps until the condensates touched and fusion started which was determined by observation of a rapid drop in the force signal (Fig. S3c). During fusion, the trap distance was then held constant. Fusion was further monitored by brightfield microscopy. Events in which fusion was not complete after 30 s were counted as non-fusing and assigned a fusion velocity of 0.

Analysis was performed using custom written code for the IGOR Pro 8 software (WaveMetrics, USA). Fusion velocity was determined by fitting a generalized logistic function (Eq. 5) to the differential force data along the x-axis. With F(start) and F(finish) the differential forces before fusion onset and after finished fusion, respectively.

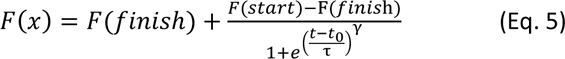

The force was then normalized so that F(start) = 0 and F(finish) = 1. The fusion velocity was determined as the slope of the tangent fitted to the normalized force data at the inflection point of the fitted function. As the fusion velocity is inversely proportional to the size of the condensates, the obtained velocity was normalized using the mean size of the two fused condensates which was obtained using radial profiling of the brightfield videos pre-fusion. For comparison between conditions velocities were transformed into the log10-fold change relative to the mean of the chromatin only condition. Statistical significance of the velocity changes was determined by t-test. Data is visualized in boxplots showing the median ± quartiles, with whiskers indicating the 9^th^ and 91^st^ percentile.

### ThT fluorescency measurment

Thioflavin T (ThT) fluorescence intensity was measured with increasing ISWI concentrations with 90 nM 13mer in modified PSB (supplemented with 2 mM MgCl_2_, 2.5 mM DTT and 40 µM ThT).

### Molecular dynamics simulations

Molecular dynamics simulations were performed using custom written software in C++ and CUDA. Chromatin arrays were described as Rouse chains with an equilibrium distance between nucleosomes of 16 nm and a spring constant of 15 pN/nm. All nucleosome particles further interacted with each other via truncated Lennard-Jones potentials. The remodeler was modeled as single particles that repel each other and can form two independent interactions with nucleosomes (Fig. S4a). All particle interactions were described as truncated Lennard-Jones potentials with an equilibrium distance of 16 nm and a cutoff distance of 48 nm. Between 32 and 48 nm the potential was simplified using a linear approximation (Fig. S4c). The strength of the remodeler-nucleosome interactions was modulated based on the remodeler’s nucleotide state. The first interaction site switched from a remodeler off-rate of 2.5*10^-6^ timestep^-1^ in the nucleotide-free state to an off-rate of 1.25*10^-7^ timestep^-1^ in the ATP and ADP bound states. The second interaction site had a remodeler off-rate of 1.25*10^-6^ timestep^-1^ in the ADP bound state and 1.25*10^-7^ timestep^-1^ in the ATP bound state and without bound nucleotide. Condensates were formed from 169 Rouse chains of length 13 and 440 remodeler particles that were equilibrated in their respective nucleosome condition for 4*10^7^ timesteps before use in FRAP or fusion simulations. Particle motion was modeled using the Langevin equation with a diffusion coefficient of 0.02 nm^2^/timestep (8*10^5^ nm²/s) corresponding to remodeler diffusion *in vivo* (Kim *et al*., 2021) in timesteps of 20 ns.

Apart from simplifications due to coarse-graining, the prime reason why we achieve only qualitative, rather than quantitative agreement between the coarse-grained simulations and experiments lies in technical constraints required for computational performance. To keep the simulation tractable, typical simulated droplet sizes are approximately an order of magnitude smaller than their experimental counterparts. Even in the model of ISWI being a stable crosslinker, a small shape change during droplet fusion may be expected from the stretching of arrays. This relative change is much more pronounced for the smaller simulated droplets, compared to the experimental droplets. In addition, simulated fusion trajectories have a well-defined endpoint (circularity=1), such that we were able to determine a fusion velocity for all simulation trajectories. In contrast, the fusion velocity for experimental trajectories could only be determined for events which completed fusion (see previous section).

### Simulated FRAP

For the simulation of FRAP experiments, remodeler particles in one half of the condensate were marked as bleached. Mixing was determined using the distances between particles. First, pairwise distances either between all particles of one type (distall) or only between bleached and non-bleached particles (dist_cross_) were determined.

The pairwise distances were binned, and mixing (M) was quantified with Eq.6

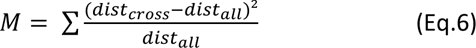

This was done at timesteps 2.5*10^6^ timesteps apart. The resulting values were normalized between 0 and 1 to emulate FRAP recovery. For each condition three simulations were performed. Data was visualized in boxplots showing the median ± quartiles and whiskers showing the maximum and minimum values.

### Simulated fusion

For the simulation of condensate fusion, two preequilibrated condensates were positioned in close proximity. Fusion progress was then measured using the sphericity of the system over time which approaches 1 as fusion progresses. The circularity was then fitted with the same generalized logistic function as the force response in the experiments. Three simulations were performed for each condition. Relative fusion velocities were obtained by normalizing the data to the mean of the nucleotide free condition. Data was visualized as boxplots as described for simulated FRAP.

## Acknowledgements

We thank Sabrina Albig for Cy3-labeled histones, Ameli Lentz for purifying GFP and cloning ISWI-GFP, Marlies Muernseer (Nanolive) for collecting holotomography data, Madhura Khare and Silvia Härtel for help with protein purification, and the Ökten group for providing sfGFP plasmid. P.V. acknowledges support from the IRTG SFB 1064. F.M.-P. acknowledges financial support from the Deutsche Forschungsgemeinschaft (SFB1064 A07, MU3613/3-1, MU3613/8-1); J.S. from the LMU Center for Nanoscience CeNS, a DFG Emmy Noether grant (No. STI673/2) and an ERC Starting Grant (No. 758124); P.B.B from SFB1064 A01 and BE1140/6-1; and M.H. by St. Jude Children’s Research Hospital, the American Lebanese Syrian Associated Charities and NIH awards R01GM141694 and R01GM135599.

## Author contributions

Conceptualization: F.M.-P., J.S., P.V.

Methodology and formal analysis: P.V., J.S., M.G.P, N.H., A.S., D.K., M.S.

Investigation: P.V., M.G.P., A.S., J.B., J.S., D.K., N.H., M.S.

Writing of original draft: F.M.-P., P.V., M.G.P, D.K., M.S., A.S., J.B., J.S.

Visualization: P.V., J.S., D.K., M.S., M.G.P.

Funding acquisition: F.M.-P., J.S., P.B.B, M.H.

Writing—Review and Editing: All authors

## Data availability statement

Authors can confirm that all relevant data are included in the article and/or its supplementary information files.

## Code availability statement

Code available on request from the authors. The MD simulation executable as well as analysis scripts for MD simulations and controlled condensate fusion experiments are available at https://github.com/StiglerLab/Vizjak_2023.

---

**Figure S1.**
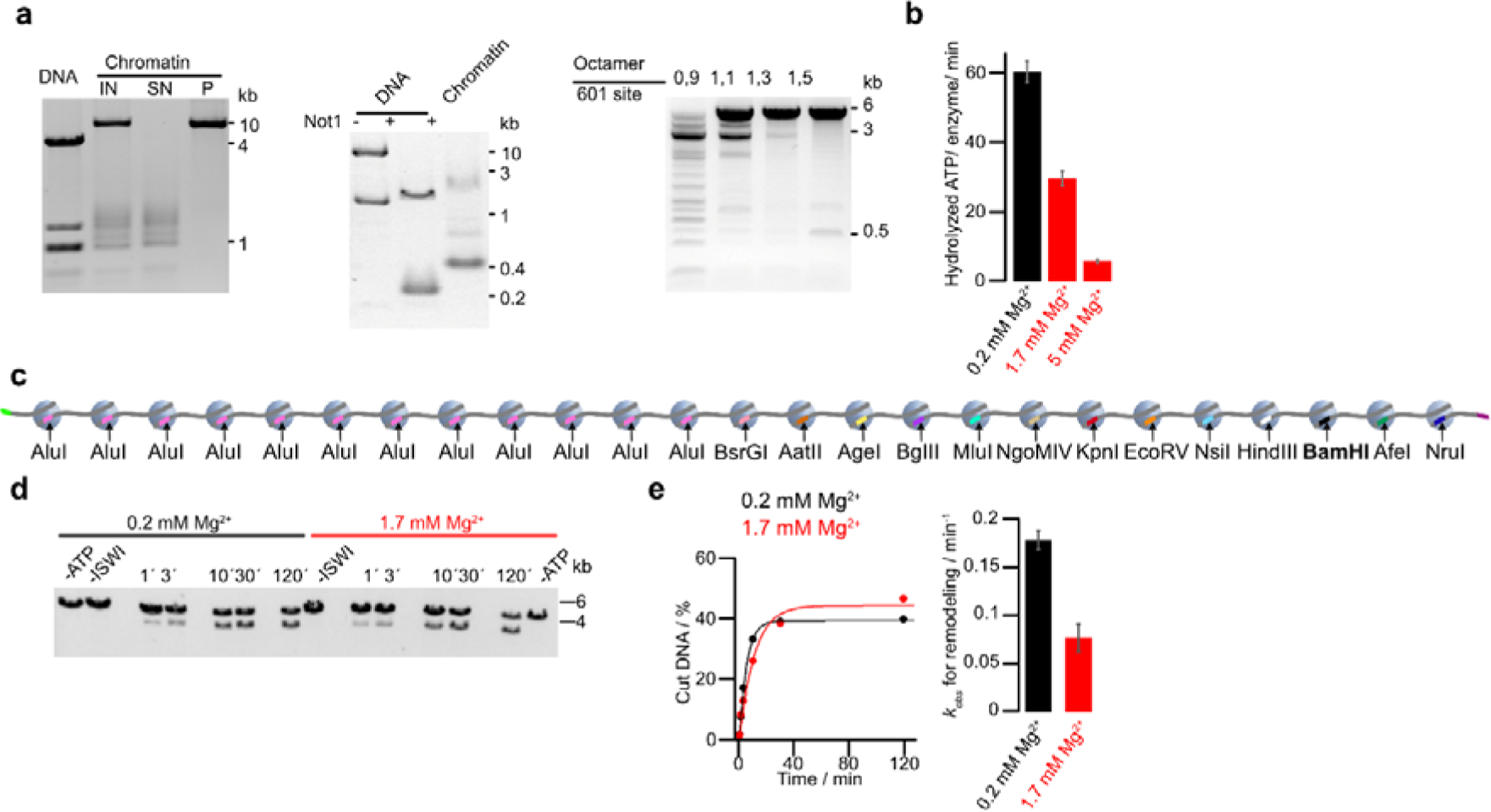
**a**, Quality controls for 25mer nucleosome arrays. Left: agarose gel after magnesium precipitation of assembled arrays and the resolubilized pellet. IN, input; SN, supernatant; P, pellet. Competitor DNA derived from the plasmid backbone (< 1 kb) was excluded from P. Middle: Not1 digestion (a Not1 site is present in each linker) liberated mostly mononucleosomes, running around 400 bp, but little 197 bp fragments, confirming saturation of most 601 repeats with octamers. Right: BsiWI digestion (all nucleosomes occlude a BsiWI restriction site) for arrays assembled with different octamer amounts. As 601 sites become saturated, digestion is hindered. **b**, ATP turnover in the presence of saturating concentrations of mononucleosomes (1.33 µM). Control experiments with three times lower mononucleosome concentrations gave the same results. Bars are mean values of two independent experiments, error bars their minimal and maximal values. Increasing Mg-concentrations reduce mononucleosome-stimulated ATP hydrolysis rates at saturating concentrations of ATP (1 mM). **c**, As in Fig. 1a, but with different orientation of the unique restriction sites such that the BamHI site is now more peripheral. **d**, BamHI accessibility assay as in Fig. 1f but for the array shown in *c*. **e**, Left: quantification of gel in *d* and exponential fits of time courses. Right: rate coefficients from single exponential fits. Bars are mean values of two independent experiments, error bars their minimal and maximal values.

**Figure S2.**
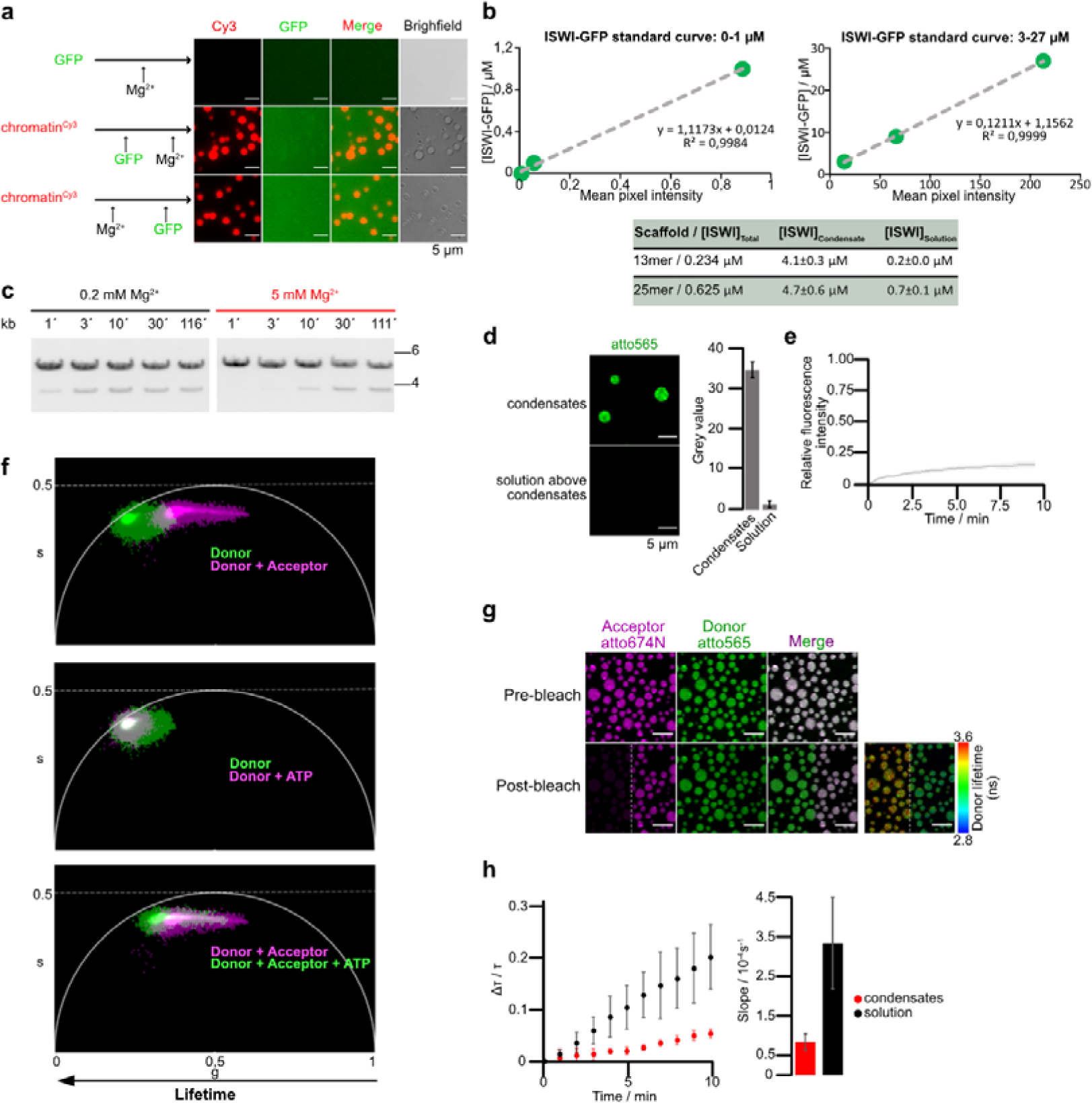
**a,** GFP does not partition into chromatin condensates. The colocalization experiment was performed with 40 nM of unlabeled 25mer, 10 nM of 25mer-Cy3 and 1.125 µM of GFP-GST. **b,** ISWI-GFP concentrations were determined inside condensates and, after centrifugation, in the surrounding solution (bottom) from fluorescence intensities using calibration curves with ISWI-GFP dilutions (top). Two different microscope settings were used to image lower and higher dilutions. Means and SD of two independent replicates are shown. **c,** Nucleosome sliding time courses in absence and presence of condensates (0.2 mM and 5 mM MgCl_2_, respectively), replicate of experiment in Fig. 2e. Sliding was measured by KpnI accessibility of 25mer arrays (15 nM). Reactions contained 750 nM ISWI and were started by addition of 1 mM Mg-ATP. **d,** Enrichment of labeled 0N60 mononucleosomes in chromatin condensates, N = 3 ± SD. The mononucleosome concentration in solution was determined from z-plane above the condensates. **e,** Whole condensate FRAP of FRET-0N60 nucleosomes to assess their exchange between condensate and solution. Line is an average and shadow SD of eight bleached condensates. One of four independent replicates with similar results is shown. **f,** Phasor representation of data in Fig. 2f. The phasor representation makes no assumptions on the number of decay rates nor on specific decay model (exponential, non-exponential) (Digman et al., 2008). Upon introduction of the acceptor, the donor’s lifetime distribution moves away from the universal circle line (single exponential lifetimes). This is consistent with an existence of at least two or even three different populations of donor: high FRET, low FRET and no FRET. **g,** Acceptor bleaching enhances donor fluorescence and lifetime, indicative of FRET. Imaging of FRET-0N60 nucleosomes in chromatin condensates. The acceptor fluorophore was bleached in the left half of the field of view, leading to an increase in donor fluorescence and donor lifetime (right most panel). **h,** FLIM-FRET measurements as in Fig. 2h, but with 5 mM Mg-ATP. Means and SD of three independent experiments.

**Figure S3.**
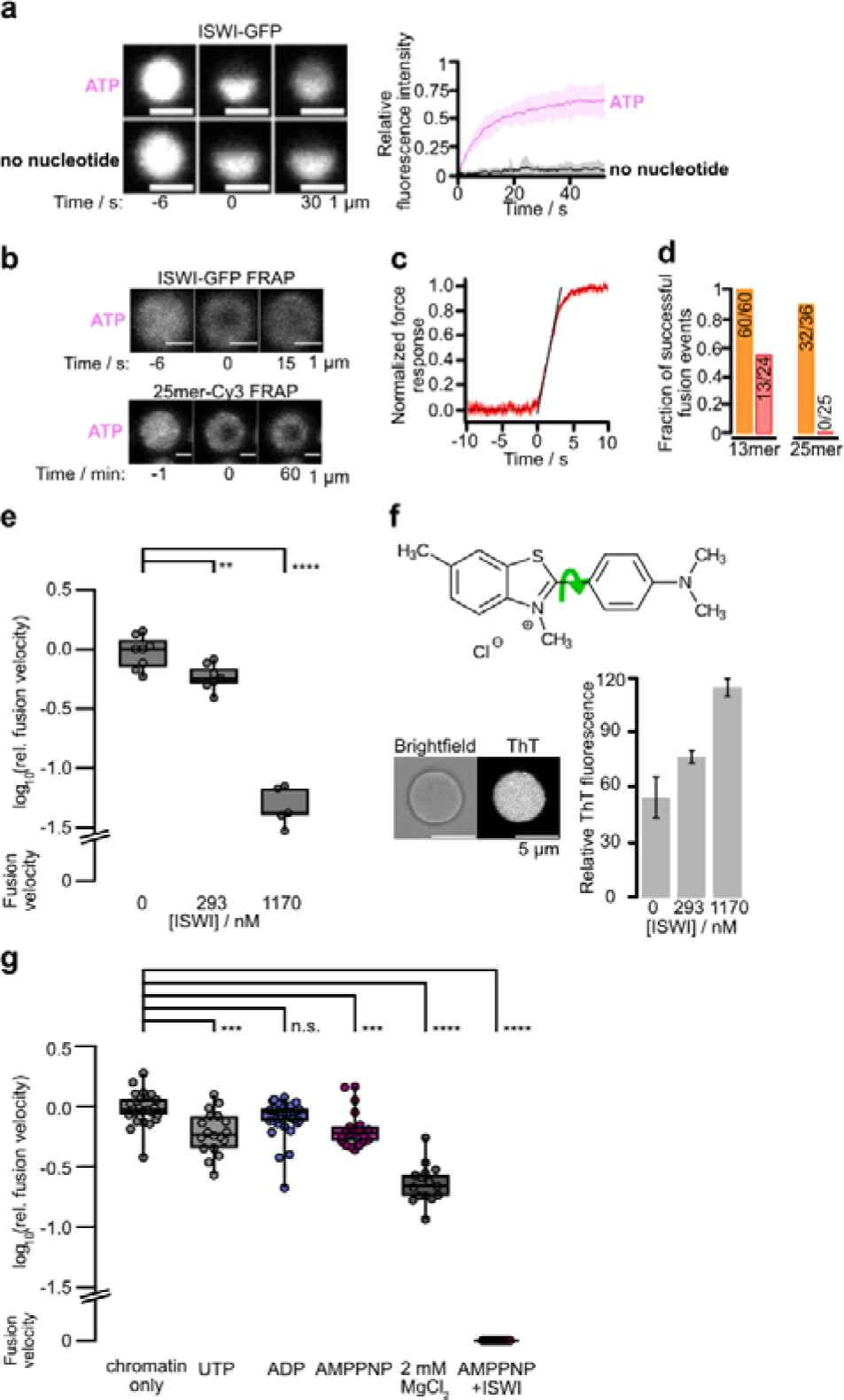
**a,** Intra-condensate mobility of ISWI-GFP measured by partial-condensate FRAP. Half of a condensate was bleached in presence and absence of Mg-ATP (1 mM). Line is an average and shadow SD of 15 bleached condensates for each condition. **b,** ISWI-GFP is more mobile than chromatin in condensates. Dual FRAP of ISWI-GFP and Cy3-labeled condensates formed by 25mer arrays in presence of Mg-ATP (1 mM) and ISWI (100 nM). Independent replicates show similar results. **c**, Force readout of optical traps during controlled condensate fusion. The fusion velocity was determined as the slope of the tangent fitted to the normalized force data at the inflection point of the fitted function. Incurred forces during fusion are on the order of 1 pN (see Methods). **d,** Fraction of successful fusion events in 1 mM MgCl_2_ (orange) and 5 mM MgCl_2_ (red). Total nucleosome concentration was 1170 nM. **e,** Fusion between condensates measured by optical tweezers containing indicated ISWI concentrations and 1170 nM nucleosomes. Velocities were estimated as illustrated in Fig. S3c. Box plots show means and 25^th^/75^th^ percentiles, whiskers the 9^th^ and 91^st^ percentiles. **f,** ISWI increased viscosity of condensates as reported by enhanced fluorescence of the molecular rotor thioflavin T (ThT). In a high viscosity medium, rotation around the C-C bond (green arrow) is constrained, and the excitation energy is released as fluorescence. Bars are average ThT fluorescence intensities relative to the outside medium; error: SD. Six condensates were analyzed for no ISWI and 293 nM ISWI, five for 1170 nM ISWI. **g**, Fusion between condensates without ISWI measured by optical tweezers. Nucleotides and free Mg^2+^ show only modest effects on fusion velocity.

**Figure S4.**
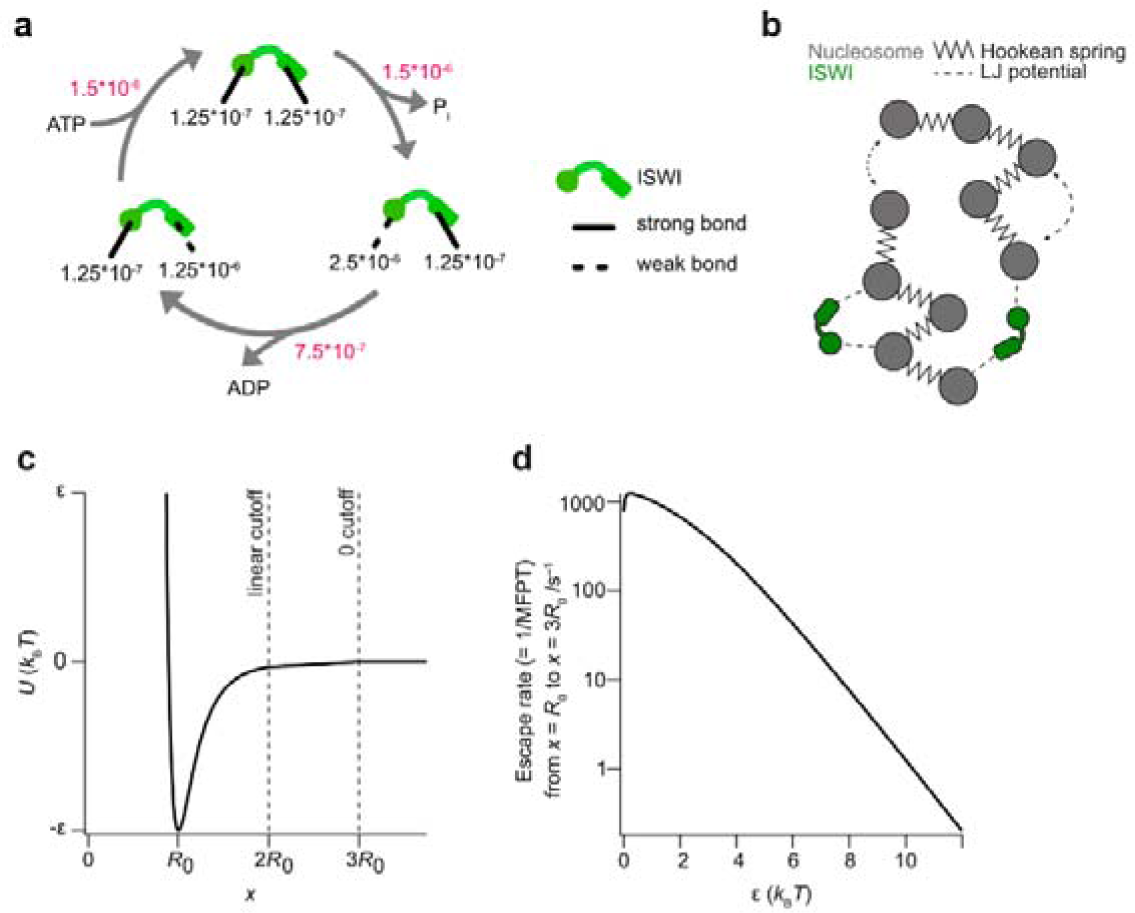
**a,** A model for the independent switching of the strengths of the two nucleosome interaction sites during ISWÍs ATPase cycle. Escape rates (black) and transition rates (red) in timestep^-1^ are indicated. **b,** Schematic representation of the implementation of the model in Fig. 4a for molecular dynamics simulations. **c,** Modified Lennard-Jones potential used in simulations. Below distances of 2R_0_ a regular Lennard-Jones potential is used. Between 2R_0_ and 3R_0_ the potential is described using a linear approximation, while interactions with range above 3R_0_ are set to 0. **d,** Conversion of the strength of the modified Lennard-Jones potential to escape rates based on the mean first passage time of potential escape (Gray & Yong, 2021).

**Movie S1. FLIM-FRET timelapse with ATP Movie S2. FLIM-FRET timelapse with AMPPNP**

**Movie S3. Controlled fusion with optical tweezers of chromatin condensates containing ISWI and ADP-BeFx**

**Movie S4. Controlled fusion with optical tweezers of chromatin condensates containing ISWI and ATP**

**Movie S5. Simulation of ISWI FRAP with graph**

**Movie S6. Simulation of condensate fusion with graph**

**Table S1.**
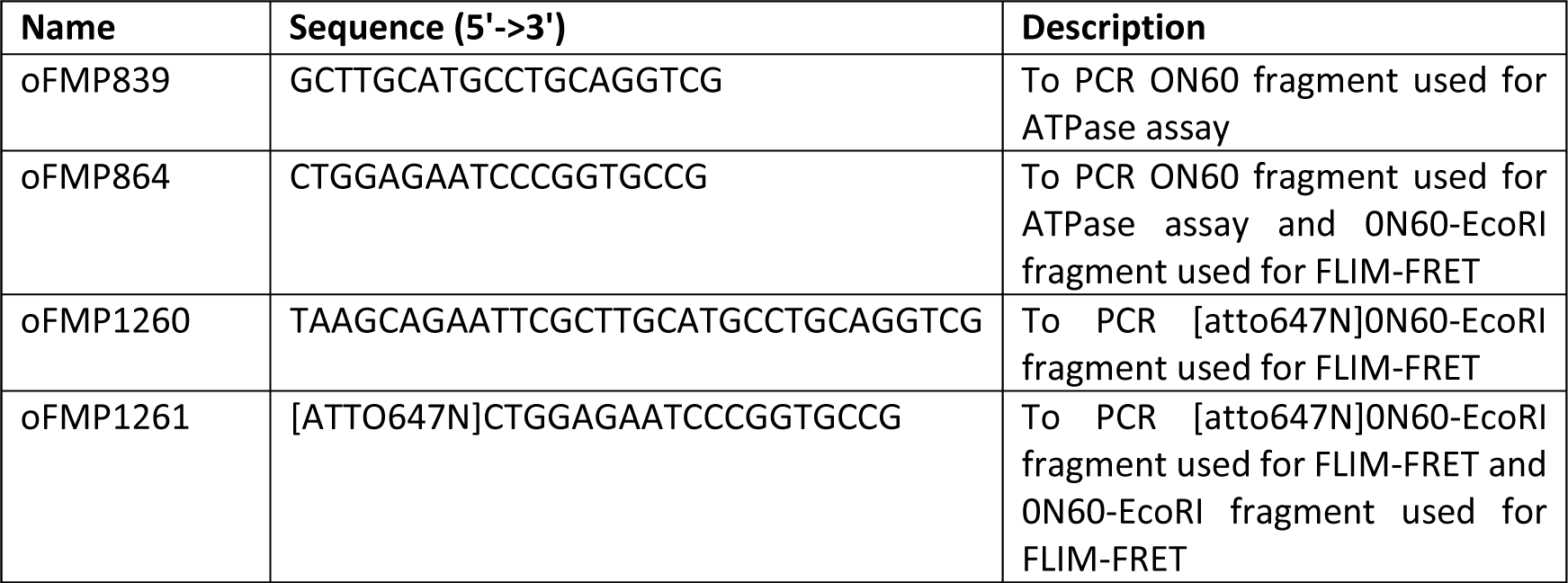
Oligonucleotide list.

